# Poly-omic risk scores predict inflammatory bowel disease diagnosis

**DOI:** 10.1101/2022.09.14.508056

**Authors:** Christopher H. Arehart, John D. Sterrett, Rosanna L. Garris, Ruth E. Quispe-Pilco, Christopher R. Gignoux, Luke M. Evans, Maggie A. Stanislawski

## Abstract

Inflammatory Bowel Disease (IBD) is characterized by complex etiology and a disrupted colonic ecosystem. We provide a framework for the analysis of multi-omic data, which we apply to study the gut ecosystem in IBD. Specifically, we train and validate models using data on the metagenome metatranscriptome, virome, and metabolome from the Human Microbiome Project 2 IBD Multi-omics Database, with 1,785 repeated samples from 131 individuals (103 cases, 27 controls). After splitting the participants into training and testing groups, we used mixed effects least absolute shrinkage and selection operator (LASSO) regression to select features for each -omic. These features, with demographic covariates, were used to generate separate single-omic prediction scores. All four single-omic scores were then combined into a final regression to assess the relative importance of the individual -omics and the predictive benefits when considered together. We identified several species, pathways, and metabolites known to be associated with IBD risk, and we explored the connections between datasets. Individually, metabolomics and viromics scores were more predictive than metagenomics or metatranscriptomics, and when all four scores were combined, we predicted disease diagnosis with a Nagelkerke’s R^2^ of 0.46 and an AUC of 0.80 [95% CI: 0.63, 0.98]. Our work suggests that some single-omic models for complex traits are more predictive than others, that incorporating multiple -omics datasets may improve prediction, and that each -omic data type provides a combination of unique and redundant information. This modeling framework can be extended to other complex traits and multi-omic datasets.

**Importance:** Complex traits are characterized by many biological and environmental factors, such that multi-omics datasets are well-positioned to help us understand their underlying etiologies. We applied a prediction framework across multiple -omics (metagenomics, metatranscriptomics, metabolomics, and viromics) from the gut ecosystem to predict inflammatory bowel disease (IBD) diagnosis. The predicted scores from our models highlighted key features and allowed us to compare the relative utility of each -omic dataset in single-omic versus multi-omics models. Our results emphasized the importance of metabolomics and viromics over metagenomics and metatranscriptomics for predicting IBD status. The greater predictive capability of metabolomics and viromics is likely because these -omics serve as markers of lifestyle factors such as diet. This study provides a modeling framework for multi-omic data, and our results show the utility of combining multiple -omic data types to disentangle complex disease etiologies and biological signatures.

## Introduction

Inflammatory Bowel Disease (IBD) is characterized by complex etiology and contains multiple pathological subtypes, including Crohn’s disease (CD) and ulcerative colitis (UC). Both of these subtypes have wide-ranging heritability estimates and involve disruptions of the gut mucosa as well as dysbiotic gut microbiota. Despite clustering of these diseases within family trees, genome-wide association studies have produced much smaller SNP-heritability estimates (0.37 for CD and 0.27 for UC)^1^ than twin studies, which estimate heritability coefficients of 0.75 for CD and 0.67 for UC.^2^ IBD affects individuals worldwide and has an incidence of 19.2 per 100,000 person-years in North America,^3^ and age-adjusted prevalence of 0.40% and 0.65% for CD and UC respectively.^4^ With impactful symptoms including diarrhea, abdominal pain, bloody stools, weight loss, and fatigue, the rising rates of CD and IBD are a salient public health concern.^5^

In genome-wide association studies to date, more than 120 related genes have been identified for CD, 67 of which were found to be differentially expressed for both CD and UC patients compared to non-IBD individuals.^6^ These biological underpinnings have connected IBD to an array of comorbidities including asthma^7^ and diabetes,^8^ which are also characterized by polygenic architectures and complex environmental interactions related to immune activation. Additionally, many of the genes most strongly associated with IBD diagnosis are active in host-microbe interactions (such as pathogen-associated molecular pattern recognition, inflammatory responses, and phagocytic processes), including toll-like receptor (TLR) 4, TLR9, nucleotide binding oligomerization domain containing 2 (NOD2), interleukin-23 receptor, and tumor necrosis factor.^6^ Multiple environmental risk factors for IBD have been identified, including smoking, excess body fat,^9^ urban living,^10^ antibiotic exposure,^10^ and adverse childhood events.^11^ On the other hand, studies have highlighted protective environmental factors including *Helicobacter pylori* infection,^12^ breastfeeding,^9^ and adequate vitamin D status.^10^ Notably, many of these environmental factors are associated with changes to an individual’s gut microbiome, its activity, the host’s metabolome, and immune function.^13,14^

Our aims for this study were two-fold: (1) develop a generalizable and interpretable modeling framework that can used to study multi-omics datasets for complex traits by evaluating the predictive contribution of each -omics data type, and (2) apply this framework to the gut ecosystem in individuals with and without IBD to contextualize important features across 4 -omics datasets. To accomplish both of these objectives, we reached beyond genetic data and genome-wide association studies to further uncover risk factors for IBD. This included microorganisms (both bacteria and viruses), RNA transcripts of the microbiome, and small molecules (metabolites) present in the gut. Whereas most studies including -omics analyses use univariate differential abundance methods (examining one feature at a time), our goal was to create an additive and predictive multivariable model that would incorporate features from multiple -omic data types. Given that many -omics data types are compositional^15,16^ (meaning that the data are proportions or relative abundances constrained by library size), a decrease in one feature will correspond to an increase in others. Thus, multivariable differential abundance analysis or models for predicting disease status based on -omics data (as will be described later in this paper) may provide a unique alternative to univariate differential abundance methods.

Analogous methods to polygenic risk scores^17^ have begun to target sources of biological variation beyond genotypes. A recent study in the UCLA Health biobank found that methylation-based risk scores were considerably more accurate than their genetic-variant-based counterpart when imputing diagnoses and lab tests in the electronic health record.^18^ This recent pairing of machine learning methods and biological data from multiple -omics levels has opened the door to better-informed models that may have greater potential for disease prevention and personalized medicine applications, especially among complex traits. The microbiome, transcriptome, virome, and metabolome, are all dynamic through time for individuals, which differs from genetic variants, which remain constant from birth. Therefore, a multi-omics-based prediction appears to be a promising approach in the context of complex and dynamic disease etiologies. Although high throughput technologies and data generation are on the rise, there are still common issues such as overfitting to training datasets, low interpretability, and the general lack of portability from one context to another.^19^ We aim to address these shortcomings by developing a generalizable and interpretable modeling framework that can be applied across the phenome and to multiple, different -omics. In the present analysis, we designed a modular multi-omics framework that allows researchers to assess the relative contributions of each -omics data type individually and then combine them together to assess overall predictive capability.

## Methods

### Data acquisition

The Human Microbiome Project 2^20^ (HMP2) IBD multi-omics database contains 1,785 unique samples collected from 131 participants across 5 study sites, with metadata including each sample’s participant ID, sex, race, antibiotic use, and site. The data used in our study were publicly available (accessed from [https://ibdmdb.org/]) and collected following approval by multiple institutional review boards, referenced in the flagship HMP2 paper.^20^ All sample collection and participant involvement was reported to follow Declaration of Helsinki guidelines and federal regulations.

### Dataset description

Each of the 1,785 samples was analyzed across up to eight different -omics data types. In this study, we used feature count tables for samples across four of the present -omics data types: metagenomics (MGN), metatranscriptomics (MTS), viromics (VRM), and metabolomics (MBL). MGN data consisted of taxonomic counts at the species level, whereas MTS data were functional profiles at the pathway level. For clarification, the term “sample” is used in this paper to denote one fecal specimen, which could have corresponding MGN, MTS, VRM, and MBL data, whereas “participant” is used to denote an individual who may have multiple samples throughout the duration of the study. MBL, MGN, MTS, and VRM data were collected on average 10.3, 3.3, 6.5, and 7.4 weeks apart per participant, respectively (see Supplementary Figure 1). Count tables for MGN and VRM were generated with MetaPhlAn2 and VirMap, respectively. MTS count tables were generated by summing pathway abundances as mapped by HUMAnN2. For MBL, four column methods (Carbon 18 (C18)-positive, C18-negative, Hydrophilic interaction liquid chromatography (HILIC)-positive, and HILIC-negative) coupled with mass spectrometry were used to isolate and detect metabolites. Only compounds successfully annotated with molecule names were included in analyses.

### Data processing and thresholding

Compositional datasets (MGN, MTS, VRM) were normalized with center log-ratio transformation, and MBL was normalized using a log_10_ transformation. We then removed highly sparse features (found in fewer than 5% of samples) from VRM. For MBL, we only included compounds present in >99% of samples. The difference in methods used for these two data types is due to variation in sparsity of the datasets, as the majority of viruses were found in very few samples, whereas the majority of compounds from MBL were found in most samples. Additionally, we removed one of any two highly collinear features (Pearson’s *ϱ* > 0.95) from MBL and MTS datasets at random. After normalization, features with a standard deviation less than 1 were excluded from MGN and MTS, and features with a standard deviation less than 0.1 were excluded from VRM and MBL. The standard deviation threshold for each -omics data type was chosen based on a histogram of sample variation in the dataset and served to eliminate features with minimal differences across samples.

### Training and validation split

Represented in our sample set were 130 distinct individuals (one participant was excluded due to missing data). To avoid overfitting, samples from 30 participants (23% of individuals) were set aside to serve as a validation set for performing model predictions and evaluating the resultant accuracy, as well as for use in our multi-omic model. Because the multi-omic model required all four data types, the 30 participants who had the most samples with all four -omics data types and the fewest samples with missing -omics were chosen for the validation set (Supplementary Figure 2). We ensured that no samples from these individuals were used in the training process for any of the four -omics layers. Any samples in the validation set with missing -omics were excluded from the multi-omic model. No aspects of the experimental design indicate bias for how many samples from a participant contain all four -omics (based on any qualities of that participant), so this is assumed to represent a non-biased subset of the participants to the best of the researchers’ knowledge from all available information on sample collection. The remaining training set contained 100 participants with a varying number of samples across layers. Of the 30 participants reserved for validation, 15 (499 samples) had CD, 3 (102 samples) had ulcerative colitis, and 12 (427 samples) were non-IBD controls. Of the 100 training participants, 50 (1,242 samples) had CD, 35 (932 samples) had UC, and 15 (520 samples) were non-IBD controls. Supplementary Table 1 shows the breakdown of training and validation samples within each -omic data type.

### Individual-omic models

Features for each -omics data type were selected by applying mixed effects least absolute shrinkage and selection operator (mixed effects LASSO)^21^ logistic regression using the glmmLasso function in the R package glmmLasso V1.5.1^22^ using the following model, where y denotes IBD diagnosis, μ denotes the intercept, β _*i*_ denotes the effect sizes, *n* denotes the number of -omic features, *(1*|*x)* indicates a random effect for variable *x*, and ϵdenotes the error:

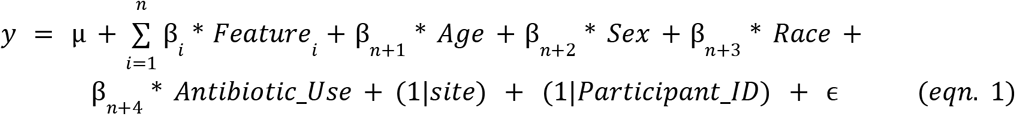

The mixed effects LASSO regression was run across a grid of lambda values, which control how parsimonious the models are (via penalizing additional features). A final LASSO regression was run using the lambda value at the elbow of the number of features retained by LASSO (Supplementary Figure 3). Using only the LASSO selected features, we generated a logistic mixed effects model (R package lme4 V1.1-29)^23^ following eqn. 1 and requiring that the important covariates of age, sex, race, antibiotic use, site, and participant ID were included. Thus, feature weights were estimated in a context that accounted for these potentially confounding variables. Analogous to the methods for computing polygenic risk scores from a genome-wide association study,^24^ the *n* feature weights (excluding participant demographic information) were then multiplied by each *i*^*th*^ samples’ feature value and summed, i.e., for sample i:

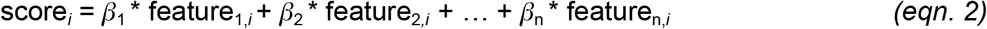

These raw scores were standardized to have a mean of zero and a standard deviation of one.

### Baseline models

We fit a baseline risk score model for each -omics data type based on only metadata for each participant using the lmer function via the lme4 R package where terms are the same as described above:

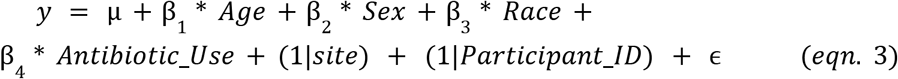

### Assessing model validity

Within each -omic data type, predicted risk scores for each individual were averaged across their longitudinal samples in the validation set (see Supplementary Figure 4). Area under the curve (AUC) values were calculated from receiver operating characteristic curves (ROC) for each model (both baseline and feature) across -omics layers. Odds ratios (ORs) were also calculated for each of the predicted scores, and 95% confidence intervals were generated for each OR.

### Multi-omic models

Prediction scores generated via feature weights within each -omics layer were averaged across samples with all 4 -omics present for each individual in the testing dataset and then combined in the following logistic regression model:

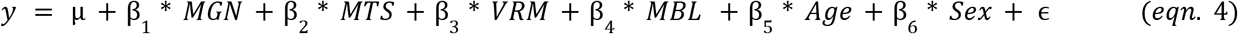

Age and sex were the two covariates included in this model due to little or no variability across the validation samples for other variables such as race or antibiotic use. Nagelkerke’s R^2^ was used to compare the combined logistic model to a baseline model only based on age and sex. Code for these methods is made publicly available at the following repository: https://github.com/sterrettJD/poly-omics-risk.

### Analyzing metabolite origins

We used Annotation of Metabolite Origins using Networks (AMON)^**25**^ to link the MBL dataset to the MGN dataset. AMON uses Kyoto Encyclopedia of Genes and Genomes (KEGG)^26^ Orthologies (KOs) present from microbial taxa and the host genome to resolve which KEGG compounds were more likely derived from microbial sources, the host, or other sources. We mapped the names of annotated, LASSO-selected compounds to KEGG compound identifiers, which involved collapsing duplicated names to single compound identifiers. A list of KOs present in the human genome was downloaded directly from KEGG, using the “hsa” organism code. The list of microbial KOs was derived from the publicly available functional profile of the MGN dataset generated using the HUMAnN pipeline.

## Results

### Included participants

This study included 130 participants, of whom, 27 did not have IBD, 65 had CD, and 38 had UC. Male and female participants were represented similarly, and the mean age of participants was 28 years old, with a standard deviation of 17 years and a range of 6 to 76 years of age. The majority of participants were white and did not use antibiotics. Table 1 elaborates with descriptive statistics of each group.

**Table 1.** Descriptive statistics of participants in the study. Categorical data are presented as “N (percent)”, and numerical data are presented as “mean (standard deviation)”. Abbreviations: SD, standard deviation, MGH, Massachusetts General Hospital.

|  | Healthy<br>Control<br>(N=27) | Crohn's<br>Disease<br>(N=65) | Ulcerative<br>Colitis<br>(N=38) | Total<br>(N=130) |
| --- | --- | --- | --- | --- |
| **Sex** |  |  |  |  |
| Female | 12 (44.4%) | 32 (49.2%) | 20 (52.6%) | 64 (49.2%) |
| Male | 15 (55.6%) | 33 (50.8%) | 18 (47.4%) | 66 (50.8%) |
| **Age** |  |  |  |  |
| Mean (SD) | 29 (± 20) | 26 (± 16) | 29 (± 17) | 28 (± 17) |
| **Antibiotics** |  |  |  |  |
| No | 27 (100%) | 51 (78.5%) | 33 (86.8%) | 111 (85.4%) |
| Yes | 0 (0%) | 14 (21.5%) | 5 (13.2%) | 19 (14.6%) |
| **Site** |  |  |  |  |
| Cedars-Sinai | 1 (3.7%) | 20 (30.8%) | 12 (31.6%) | 33 (25.4%) |
| Cincinnati | 9 (33.3%) | 17 (26.2%) | 7 (18.4%) | 33 (25.4%) |
| MGH | 13 (48.1%) | 13 (20.0%) | 11 (28.9%) | 37 (28.5%) |
| MGH Pediatrics | 3 (11.1%) | 8 (12.3%) | 5 (13.2%) | 16 (12.3%) |
| Emory | 1 (3.7%) | 7 (10.8%) | 3 (7.9%) | 11 (8.5%) |
| **Race** |  |  |  |  |
| American Indian or Alaska<br>Native | 0 (0%) | 1 (1.5%) | 0 (0%) | 1 (0.8%) |
| Black or African American | 1 (3.7%) | 3 (4.6%) | 6 (15.8%) | 10 (7.7%) |
| More than one race | 1 (3.7%) | 3 (4.6%) | 1 (2.6%) | 5 (3.8%) |
| White | 25 (92.6%) | 56 (86.2%) | 29 (76.3%) | 110 (84.6%) |
| Other | 0 (0%) | 2 (3.1%) | 2 (5.3%) | 4 (3.1%) |

### Baseline models

Baseline models, which only include patient demographic information and were trained on the training dataset, poorly predicted the actual diagnosis of our validation dataset, as can be seen in Supplementary Table 2 (MGN AUC [95% CI] = 0.43 [0.19, 0.67], MGN Nagelkerke’s R^2^ = 0.08; VRM AUC [95% CI] = 0.47 [0.23, 0.71], VRM Nagelkerke’s R^2^ = 0.02; MTS AUC [95% CI] = 0.38 [0.15, 0.62], MTS Nagelkerke’s R^2^ = 0.04; MBL AUC [95% CI] = 0.47 [0.25, 0.69], MBL Nagelkerke’s R^2^ = 0.01). This established that diagnosis could not be discriminated solely by participant metadata and non-omics covariates as described in equation 3 and Supplementary Table 1.

### Individual -omic risk scores

Compared to the baseline models, the predicted risk scores (using feature weights derived from the training set) for each -omic data type (in addition to the baseline covariates of age and sex) demonstrated better predictive capability on the validation data as is shown in Figure 1 (MGN AUC [95% CI] = 0.66 [0.44, 0.87], MGN Nagelkerke’s R^2^ = 0.12; VRM AUC [95% CI] = 0.83 [0.68, 0.98], VIR Nagelkerke’s R^2^ = 0.43; MTS AUC [95% CI] = 0.73 [0.53, 0.92], MTS Nagelkerke’s R^2^ = 0.20; MBL AUC [95% CI] = 0.82 [0.66, 0.98], MBL Nagelkerke’s R^2^ = 0.40). A similar plot showing AUCs and ORs when only using the -omic-derived scores to predict diagnosis is shown in Supplementary Figure 5 with similar results, suggesting that the covariates of age and sex are not driving this predictive accuracy (MGN AUC [95% CI] = 0.70 [0.50, 0.91], MGN Nagelkerke’s R^2^ = 0.04; VRM AUC [95% CI] = 0.80 [0.63, 0.96], VRM Nagelkerke’s R^2^ = 0.31; MTS AUC [95% CI] = 0.64 [0.42, 0.86], MTS Nagelkerke’s R^2^ = 0.12; MBL AUC [9%] CI = 0.78 [0.59, 0.98], MBL Nagelkerke’s R^2^ = 0.32). Of the individual -omics models, VRM and MBL had the best predictive capability with little to no interquartile overlap between cases and controls. The VRM and MBL predicted scores had significant odds ratios of 9.28 [1.53, 56.13] (p = 0.02) and 4.30 [1.32, 13.98] (*p* = 0.02) respectively, while MTS had an odds ratio of 2.89 [0.62, 13.52] (p = 0.2), and MGN had the lowest odds ratio of 1.24 [0.50, 3.09] (p =0.6).

**Figure 1.**
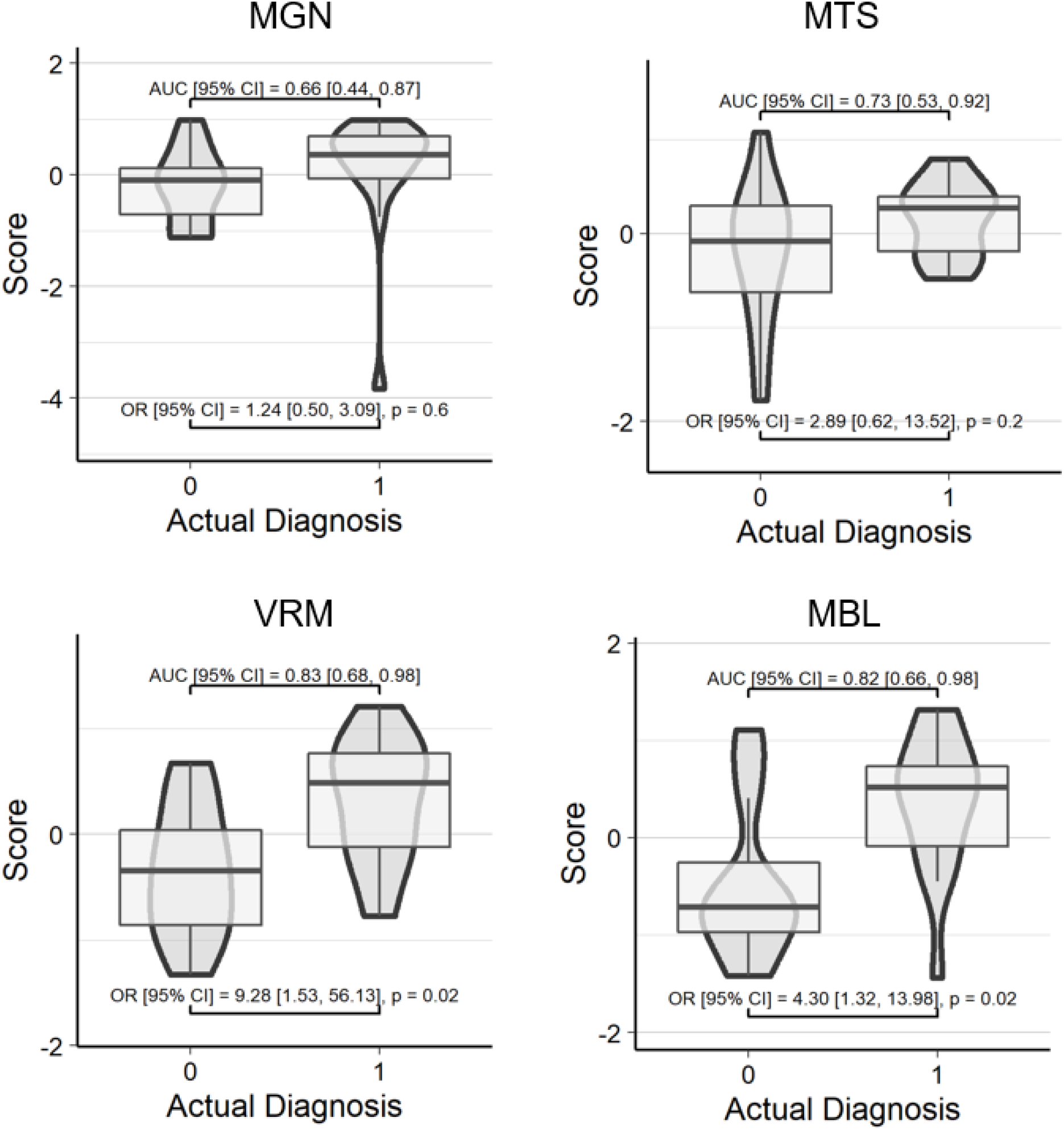
Risk scores predict IBD diagnosis. Z-score transformed risk scores (averaged across all samples for each participant) on the y axis are plotted against actual diagnosis on the x axis for the validation dataset. AUC and OR were calculated with basic covariates (diagnosis ∼ score + age + sex). Each of the four scores shown were calculated using feature weights from a LASSO-identified mixed effect logistic regression trained in a separate set of samples/individuals: diagnosis ∼ features + age + sex + race + antibiotic use + (1|site) + (1|participant ID). An actual diagnosis value of 1 indicates presence of IBD. Abbreviations: MGN, metagenomics; MTS, metatranscriptomics; VRM, viromics; MBL, metabolomics; AUC, area under the receiver operating characteristic curve; OR, odds ratio; CI, confidence interval; IBD, inflammatory bowel disease.

### Selected features

Within each individual -omic model, LASSO selected 14 species from the 237 considered in the MGN dataset, representing 5.91% of all features passing quality control (QC); 23 pathways from the 280 considered in the MTS, representing 8.21% of all features passing QC, 14 features from 596 metabolites in the MTB representing 2.35% of all features passing QC, and 6 viruses from 9 VRM, representing 66.67% of all features passing QC. Figure 2 shows the feature weights for each -omics data type. For MGN, *Megasphaera* sp DISK18 had the strongest negative weight (indicating association with low risk score). Additional species associated with lower risk scores included *Parabacteroides goldsteinii, Methanobrevibacter smithii, Roseburia hominis*, and *Akkermansia muciniphila. Firmicutes* CAG 83 was the only species identified with a positive weight. Supplementary Figures 6-19 illustrate the dynamic longitudinal trends among the 14 LASSO-selected metagenomics species for the 30 individuals reserved for model validation. The variation and volatility within individuals over the 52-week sampling period highlight the importance of longitudinal sampling when trying to capture robust estimates of the effects of each taxon. Additionally, a phylogenetic tree of the MGN features shows *Alistipes putredinis* and *Alistipes shahii* are the most related species, while *Methanobrevibacter smithii* appeared to be more distantly related to the rest of the features (Supplementary Figure 20). A co-occurrence analysis revealed that the 14 selected features showed low levels of co-occurrence (<0.5 Pearson correlation index). Among the 14 features, the pairs *Eubacterium siraeum* and *Oscillibacter sp. CAG:241*, as well as *Alistipes putredinis* and *Alistipes shahii*, had the highest Pearson index of 0.5. *Roserburia hominis* and *Megasphaera sp*. DISK_18 had the lowest Pearson index of -0.03 (Supplementary figure 21-22). The absence of large negative values across the heatmap suggests that these 14 features do not have mutually exclusive relationships.

**Figure 2.**
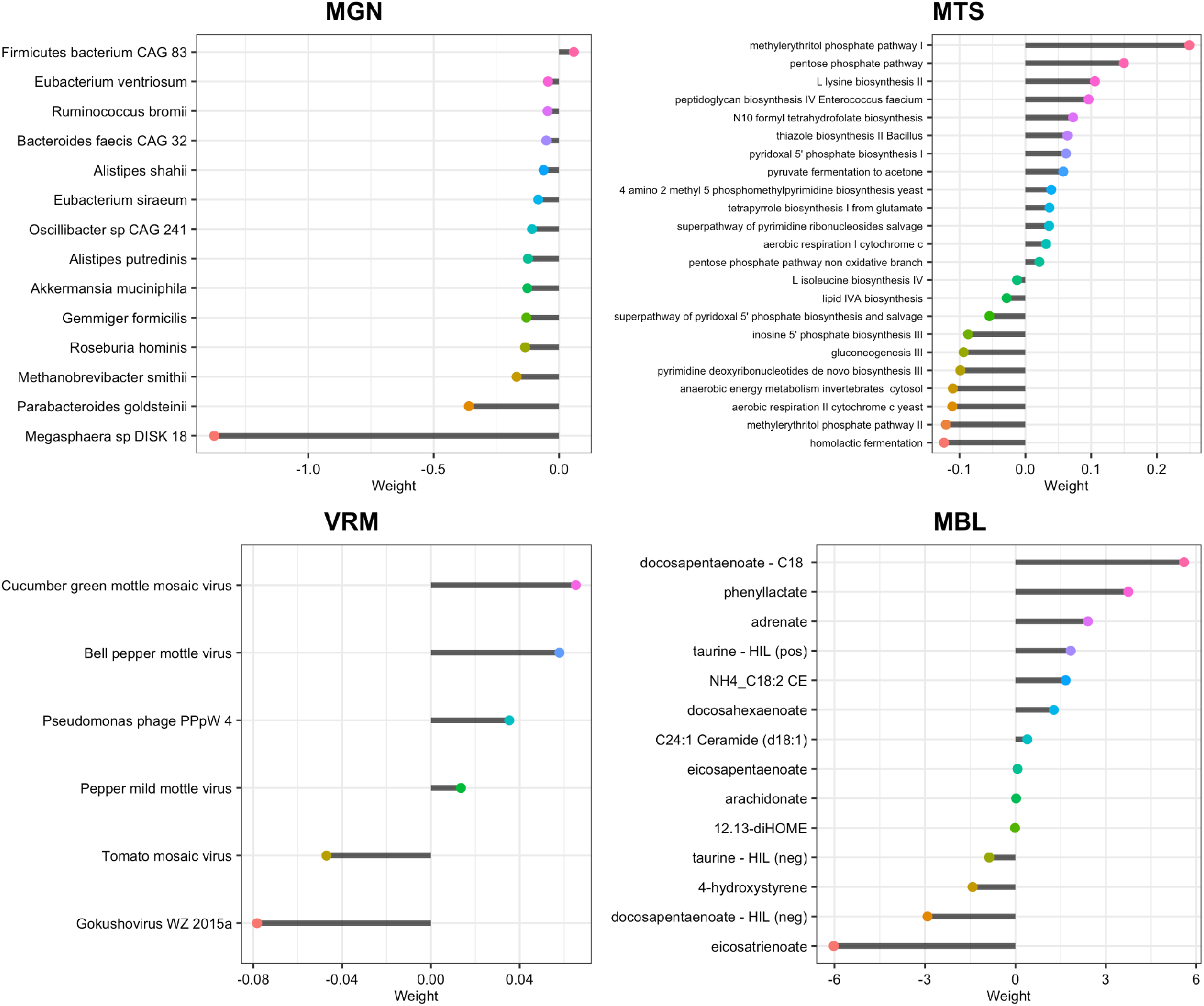
Feature weights for each risk score model. The effect sizes described by the x-axis were multiplied by transformed abundances and then summed to generate each -omic’s score. A negative weight corresponds to lower predicted risk of IBD, whereas a positive weight would confer higher predicted risk of IBD. Abbreviations: MGN, metagenomics; MTS, metatranscriptomics; VRM, viromics; MBL, metabolomics; IBD, inflammatory bowel disease; HIL, hydrophilic interaction liquid chromatography.

The MTS model selected 22 pathways. Of these, the strongest negative weights (associated with a lower risk score) were for the 2-methyl citrate cycle I and the superpathway of sulfur amino acid biosynthesis identified from *Saccharomyces cerevisiae*, and the strongest negative weights were for the pentose phosphate pathway and the methylerythritol phosphate pathway I. Of the six viruses included in the VRM model, four had positive weights indicating a direct association with IBD. The strongest positive weight was for cucumber green mottle mosaic virus and the strongest negative weight was a gokushovirus. The MBL model selected 14 metabolites. The strongest positive weights were for docosapentaenoate (identified by the C18 column) and phenyllactate and the strongest negative weights were for eicosatrienoate and docosapentaenoate (identified by the HILIC column).

### Combined multi-omics model

Figure 3a illustrates the odds ratios of all -omics scores in a combined framework for the 30 individuals reserved for model validation. The covariates of age and sex were only slightly predictive of IBD on their own (Nagelkerke Pseudo R^2^ = 0.11); however, a multiple regression including age, sex, and all four scores produced a Nagelkerke R^2^= 0.46 and an AUC of 0.80 (Figure 3c). The variance explained by the four scores without age and sex was slightly lower (R^2^= 0.37), showing the importance of including demographic covariates and the predictive capacity of our combined framework to explain variance among IBD diagnoses. Despite the 95% confidence interval for all four scores’ ORs overlapping with one, VRM and MBL stood out with larger values than MGN or MTS. The correlation between scores illustrates some similarity between VRM, MGN, and MBL, in addition to MTS and MBL (Figure 3b). However, these correlations are modest in magnitude (less than 0.5), suggesting that these scores each capture a mostly distinct signal. Analysis for a leave-one–omic-out approach is shown in Supplementary Figure 23 to further demonstrate the increased odds ratios (near the threshold for significance) for VRM and MBL compared to MGN and MTS.

**Figure 3.**
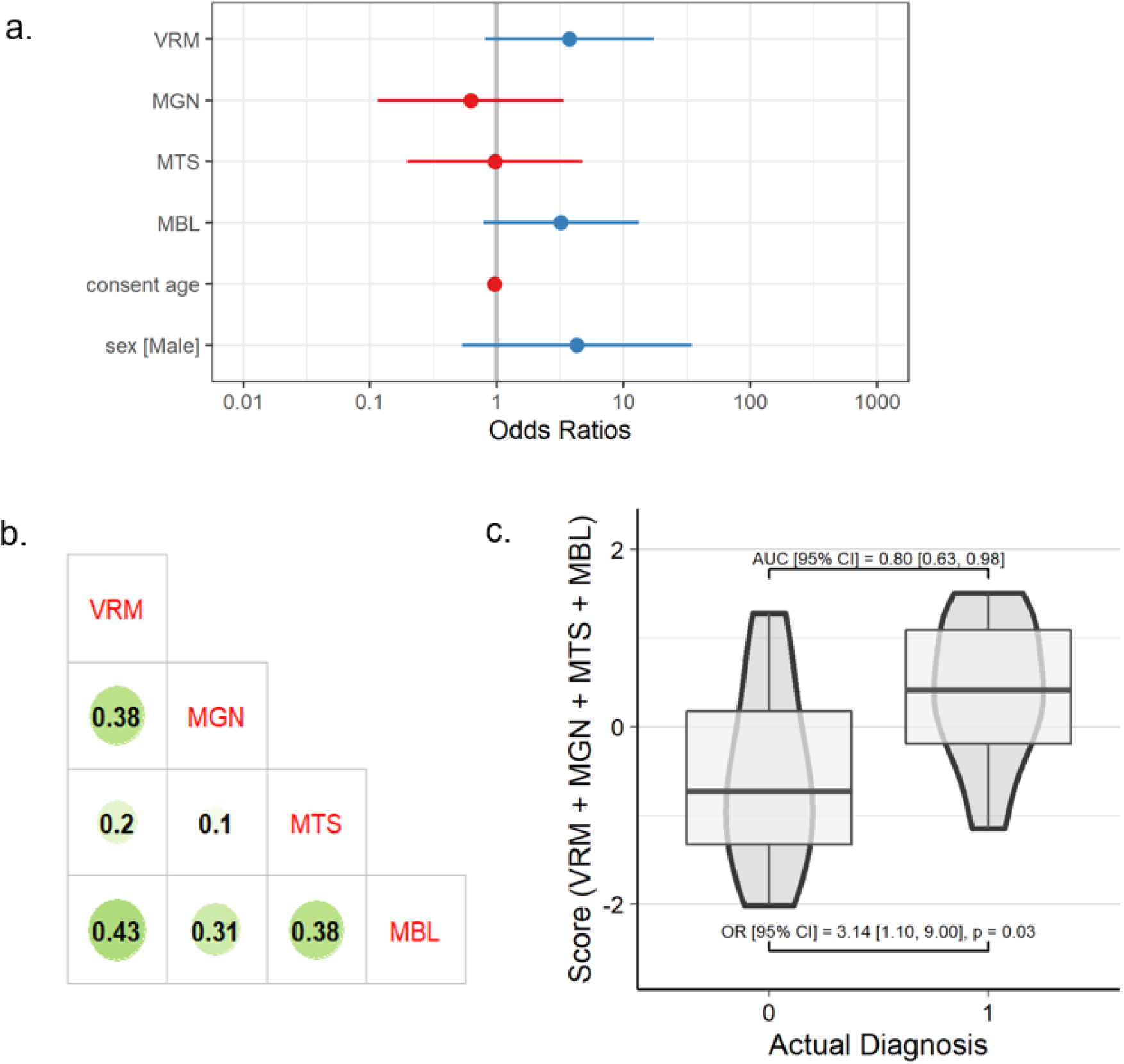
Results of multi-omic modeling. (a) Odds ratios and 95% confidence intervals of the predicted score (b) pearson correlation matrix between -omics scores, and (c) standardized risk scores (summed across all -omics) on the y-axis plotted against actual diagnosis on the x-axis for the validation dataset, where AUC and OR were calculated with basic covariates (diagnosis ∼ score + age + sex). In panel a, points represent the odds ratio for each -omic’s predicted scores in the multi-omic regression, and lines represent 95% confidence intervals of the odds ratios. In panel b, the size and darkness of the circle represents the correlation between the predicted scores for each -omic data type. In panel c, an actual diagnosis of 1 represents a case for IBD. Abbreviations: MGN, metagenomics; MTS, metatranscriptomics; VRM, viromics; MBL, metabolomics; AUC, area under the receiver operating characteristic curve; OR, odds ratio; CI, confidence interval; IBD, inflammatory bowel disease.

### Metabolite origins

AMON^25^ was used to predict the origins of the 14 metabolites selected by LASSO using the host (human) genome and the genomes of the 237 bacteria species in the MGN dataset. Of the 14 selected metabolites, 2 had the same compound identifier, resulting in 12 compounds that were unique and annotated. We used a list of KOs in the human genome, and from the KOs identified by the shotgun metagenome functional profile. AMON predicted that 7 of the 12 MBL compounds were produced by either the human or the gut microbiome, meaning that 5 of the compounds were likely produced by other sources (such as plant compounds derived from dietary sources). Figure 4 shows the classification of these 7 compounds based on their likely organismal source. Of the 7 identified compounds, 4 were likely only produced by the human, 2 were produced by either the human or the microbes, and 1 was likely only produced by the microbes present in the gut microbiome. The LASSO-selected taxa contained KOs to produce 2 of the metabolites, taurine and 4-hydroxystyrene. Notably, only the LASSO-selected bacteria were capable of producing 4-hydroxystyrene, whereas the non-selected taxa and host were not.

**Figure 4.**
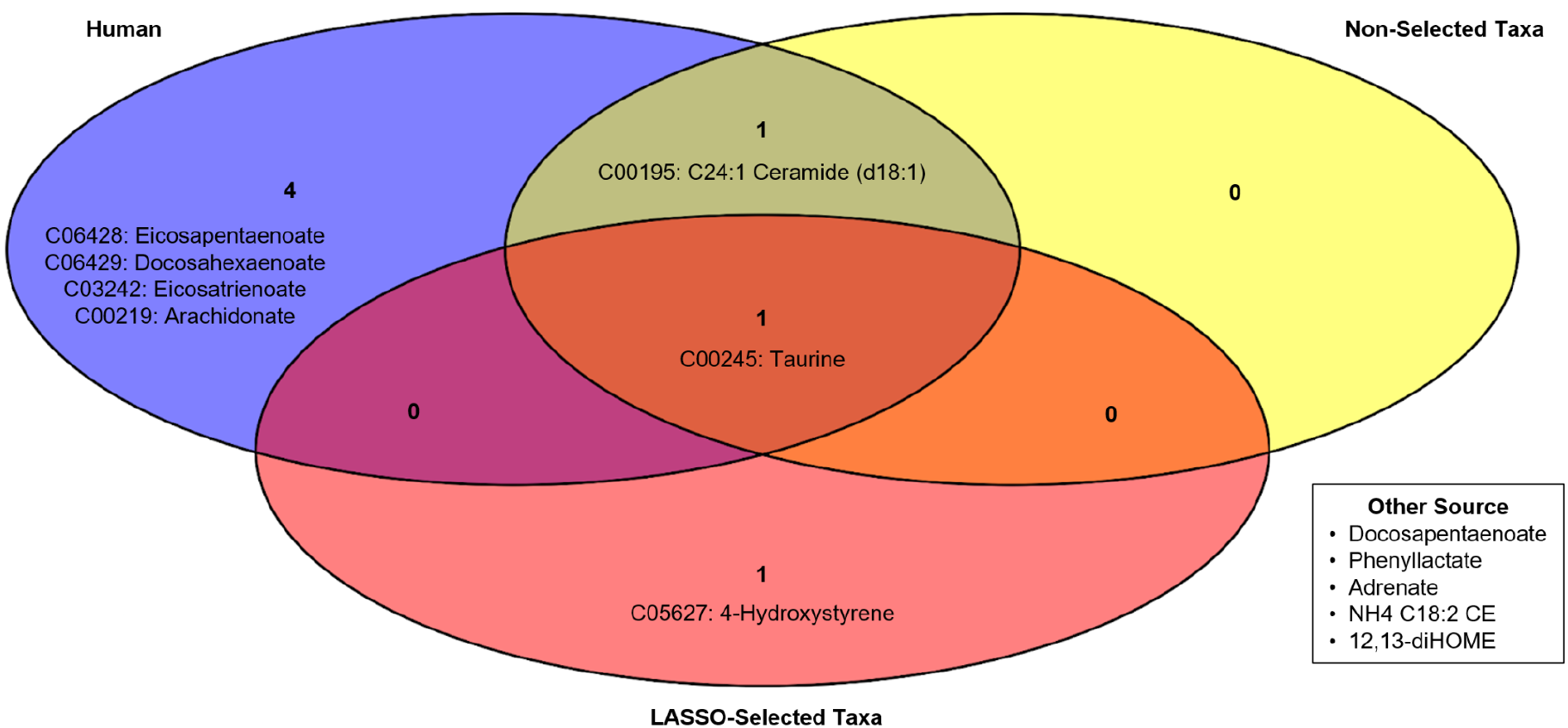
Annotation of LASSO-selected compound origins. A Venn diagram showing the likely origins of LASSO-selected compounds from the MBL dataset. “LASSO-Selected Taxa” refers to KOs from the 14 taxa selected by the MGN LASSO model, whereas “Non-Selected Taxa” refers to all KOs detected in the MGN dataset from the 223 taxa not selected by the MGN LASSO model. “Human” refers to all KOs present in the human genome, according to KEGG’s database, and “Other Sources” refers to compounds that were likely not produced by the host or detected taxa from the gut microbiome. Abbreviations: KEGG, Kyoto Encyclopedia of Genes and Genomes; KO, Kyoto Encyclopedia of Genes and Genomes Orthology; LASSO, least absolute shrinkage and selection operator; MGN, metagenomics; MBL, metabolomics.

## Discussion

Overall, we successfully utilized a polygenic risk score framework across multiple -omics data types to predict IBD diagnosis. Other studies have taken similar approaches, such as single variable differential abundance testing across multiple -omics in IBD,^27^ multivariate analysis of multi-omic interactions in participants with IBD,^28^ like a composite of unsupervised multivariate analysis and principal component analysis, or multi-omic risk scores for diseases other than IBD.^29^ Our results often corroborate the findings of other multi-omic studies of individuals with IBD, and they also highlight groups of features that associate with IBD when considering the abundances of other features. Additionally, our methods allow for comparison of the predictive ability of different -omics for disease, highlighting that taxonomic profiling (from MGN) might not hold as much of a signature of IBD as other -omics that provide information on active transcription (MTS) and compound presence (MBL). Additionally, though VRM-predicted IBD scores correlated with MGN, VIR held a much stronger relationship with IBD, which could be a result of agricultural viruses serving as markers of diet.

### Interpretation of features identified by individual models

#### Metagenomics

Among the 14 MGN features selected by LASSO, all but one had a negative weight in the multiple regression. These negative effect estimates correspond to reduced risk of IBD when utilized in the scoring framework, and the estimate with the largest magnitude (−1.37) was *Megasphaera sp. DISK 18* which is known for being an early colonizer of the oral microbiome.^30^ The next most negative effect estimate, from *Parabacteroides goldsteinii*, is of key interest given recent findings in mouse experiments that a *P. goldsteinii* probiotic may be useful in treating diet-induced obesity and type 2 diabetes.^31^ The links between the microbiome, diet, and IBD are important to consider in the context in which environmental exposures may manifest a genetic predisposition to the onset of IBD.^8^ Our findings of *Roseburia hominis* and *Bacteroides faecis CAG 32* associating with decreased risk of IBD is consistent with existing literature.^32^ We are intrigued by the importance of two features belonging to the *Alistipes* genus given the emerging connections between *Alistipes* and gut dysbiosis.^33^ Lastly, *Ruminococcus bromii*’s role in the metagenomics score is worth highlighting since *R. bromii* is known to support the growth of *R. gnavus*,^34^ which is a species purported to be associated with increased risk of CD.^32,35^

#### Metatranscriptomics

The MTS model identified a strong negative weight for transcripts belonging to the 2-methyl citrate cycle, meaning this pathway was associated with a lower risk of IBD. Notably, the 2-methyl citrate cycle is responsible for metabolizing propionic acid^36^ (a short chain fatty acid produced by gut commensal organisms fermenting dietary fiber) into succinate, which can enter the tricarboxylic acid (TCA) cycle. Increased transcription of this pathway could be indicative of high propionic acid in the gut, which has previously been demonstrated to be protective in the context of IBD, as propionic acid has anti-inflammatory effects through the inhibition of nuclear factor κB (NF-κB).^37^ Alternatively, lower propionic acid could reflect decreased dietary fiber in participants with IBD. This result also aligns with the previous analysis^38^ of the HMP2 dataset (using different methods). The MTS model additionally identified a strong negative weight for sulfur amino acid biosynthesis transcripts. Given that one signature of IBD is a microbial shift toward the catabolism of taurine and cysteine to produce hydrogen sulfide, this is consistent with previous literature as well.^39^

The MTS model identified the pentose phosphate and methylerythritol phosphate pathways as having the strongest positive weights. The pentose phosphate pathway is involved in the metabolism of C5 sugars, but not much literature exists explaining a connection between it and IBD, though one study identified an increase in the pentose phosphate pathway in ileal Crohn’s disease.^40^ Similarly, little literature exists about the potential role of the methylerythritol phosphate pathway in the context of IBD, though its intermediates are potent activators of human gamma delta T cells, which are the first line of mucosal defense.^41^ Gamma delta T cells have been implicated in intestinal inflammation and IBD in both human and animal models across a multitude of studies, though it is unclear whether their effects are protective or not.^42^

#### Viromics

The VRM model identified six viruses associated (either positively or negatively) with IBD risk. Four of these, Cucumber green mottle mosaic virus, bell pepper mottle virus, pepper mild mottle virus and tomato mosaic virus are all in the *Tobamovirus* genus. These are highly persistent and transmissible positive-sense single stranded RNA viruses that are known to infect various crop species and are found worldwide. Though many crops have developed resistance to the damaging effects of these viruses they are still endemic in agricultural products and have been found to make up a large portion of the gut virome.^43,44^ As such, presence of these viruses could be serving as a marker of food choice in these participants, given that individuals with IBD may choose to consume different foods due to their condition. Interestingly, only one of these tobamoviruses, tomato mosaic virus, had a negative feature weight. The other two viruses identified, pseudomonas phage PPpW4 and gokushovirus WV 2015a are both bacteriophages. Pseudomonas phage PPpW4 parasitizes pseudomonas bacteria, which are associated with pneumonia and post-surgical infections, and have been studied for use in phage therapy.^45^ Gokushoviruses, though ubiquitous throughout the environment, are largely uncharacterized.^46^

#### Metabolomics

The MBL model identified fourteen metabolites associated with IBD, the majority of which were fatty acids. Metabolites with positive weights were docosahexaenoate, NH_4_-18:2 cholesterol ester, C24:1-ceramide (d18:1), adrenate, phenyllactate, eicosapenatenoate, and arachidonate. Adrenate and phenyllactate have been found in increased concentrations in individuals with IBD.^20^ Metabolites with negative weights were hydroxystyrene, eicosatrienoate, and 12,13-dihydroxyoctadec-9-enoic acid (DiHOME). 12,13-diHOME was also reported as an important metabolite for differentiating IBD status when applying a knockoff filtering-based multivariate approach to data from HMP2.^47^

Our model selected two compounds annotated as taurine that had been isolated using different chromatographic columns (hydrophilic interaction liquid chromatography (HILIC) negative and HILIC positive), and interestingly, these compounds had contrasting positive and negative weights. This may be related to the charge (or other aspects) of the identified compounds, but a more detailed investigation to follow up the untargeted metabolomics would be needed to differentiate these two. In accordance with this uncertainty, multiple studies have found conflicting associations between taurine in the gut metabolome and IBD status.^48^ Taurine, being a sulfur-containing amino acid, could have very diverse roles in the gut metabolome given its microbial metabolism to hydrogen sulfide, which can have effects ranging from pathogen inhibition at moderate concentrations to direct irritation of the gut mucosa at high concentrations.^49^ Additionally, taurine plays other roles in the gut metabolome, such as conjugating with cholic acid to form the bioactive secondary bile acid taurocholic acid. Our findings are consistent with some differential abundance testing results from the original HMP2 paper, as they found taurine and taurocholic acid to be differentially abundant across IBD status.^20^

Similarly, our model selected two compounds annotated as docosapentaenoate, though in this case one with a positive weight came from the C18 column (specialized for intermediate polarity compounds, such as free fatty acids), and one with a negative weight came from the negative ion mode hydrophilic interaction liquid chromatography (HILIC) negative mode (specialized for polar compounds). According to previous studies, Docosapentaenoate is an omega-3 fatty acid generally regarded to have anti-inflammatory effects.^50^

### Comparing individual -omics models

Of the individual -omic models, MBL and VRM had the highest predictive capability, while MTS provided moderate prediction, and MGN provided the lowest predictive accuracy. One explanation for these results is that MBL and VRM provide information about both the microbiome and the host; the metabolites could have been produced by the host or their diet, and the viruses could also be diet-related given that four of the six selected viruses belong to the genus *Tobamovirus* and infect crops. In contrast, MTS and MGN only provide information about the microbiome, not the host. However, MTS specifies data on active transcription, whereas MGN only provides information about taxonomy and functional capabilities. Notably, our MGN analysis only considered taxonomy, not functional potential (i.e., no MGN pathway abundance data). These results are consistent with other findings that the metabolome predicts phenotypes better than taxonomic profiles do.^51^

### Comparison of features selected by single-omic models

AMON identified that most of the LASSO-selected compounds from the MBL dataset were not produced by the taxa selected by the MGN LASSO model. This suggests that these single-omic models are providing orthogonal information, which is complemented by the weak correlation between MGN and MBL single-omic scores. Of note, the compound 4-hydroxystyrene was identified to be produced by the MGN LASSO-selected taxa (but not the host or non-selected taxa). This has been demonstrated experimentally, as other groups have shown that microbial metabolism of hydroxycinnamic acids produces 4-hydroxystyrene in the rat microbiome.^52^ Particularly, polyphenols such as hydroxycinnamic acids (and their metabolites) have been studied as potential interventions in IBD,^53^ as they modulate epithelial inflammation and severity of Dextran Sulfate Sodium (DSS)-induced UC in mice.^54^ Thus, the integrated results of our multi-omic modeling highlight the importance of microbial metabolism of select polyphenols on IBD status and the relevance of a multi-omic modeling approach for this disease. However, with a larger sample size, an approach allowing for interactions between -omics datasets may improve the detection of microbes involved in the metabolism of phenotype-modulating compounds.

### Combined model

The combined regression model incorporating multiple -omics scores allows us to contextualize the predictive capability of each -omics model in a multi-omics context. The results of our combined model mirror the AUCs and ORs of the individual -omics models. In the combined model, MGN had the lowest OR, whereas MTS had a moderate OR, and MBL and VRM had the highest ORs. We were limited by our sample size to not be able to reserve a second bonafide validation set for the combined model, but the improved R^2^ from a baseline model indicates the predictive capability of this combined model. Other studies performing multi-omics predictive modeling should consider similar frameworks with single-omic predictive scores to improve the interpretability of important features and allow for the comparison of predictive accuracy between various -omics data types. This modular approach of building scores within each -omics data type allows for the selection of a sparse set of features within each dataset; it also leads to a combined model with insights contextualized by multiple biological -omic layers (ideally ranging from the genome to the metabolome).

### Limitations

As with many studies of this kind, batch and site effects may affect the generalizability of our model. High throughput -omics technology faces many limitations due to batch effects, and future work should consider techniques for combining batches in this framework, using batch correction techniques or incorporating random effects for batches.^55^ Such work to combine data from different studies could increase the robustness of this framework and increase the generalizability of its diagnostic capability, which would increase the likelihood of clinical implementation for such diagnostic -omic modeling, which should be considered an end goal.

Additionally, the number of -omics features was far greater than the number of participants, though we were able to use multiple longitudinal samples per participant (accounted for via random effects for participant ID). We took steps to reduce the number of features, such as through the sparsity, collinearity, and variance filtering steps, as well as through the use of LASSO regression for feature selection. However, such steps could be causing us to unwittingly discard useful information. Importantly, our modeling did not consider interactions between features, either within or between -omics, which is a direction for future research. Moreover, the use of a center log-ratio transformation is not robust to shifting total microbial biomass, nor is it subcompositionally consistent. Other log-ratio transformations, such as the isometric log-ratio transformation, are subcompositionally consistent, though they do suffer from more difficult interpretability. Future work should consider the use of additive, isometric, or phylogenetic isometric log-ratio transformations to transform compositional data out of the simplex into Euclidean space, though interpretability and use cases of poly-omic models should be considered as a tradeoff.

The relatively small sample size and sample overlap across -omic datasets limited our ability to detect weak effect sizes across features. Extending these analyses to larger and more diverse studies would allow us to evaluate these methods on a larger scale. For example, performing cross-validation splits of the training and testing datasets would have further tested the robustness of these results. However, without sufficient sample sizes overlapping between -omics layers we were limited to one training-testing split of the data. In addition, exploring non-linear machine learning techniques such as random forests may have yielded better results and motivates an exciting direction for future multi-omic analyses.

The currently available dataset from HMP2 does not allow public access of host genome data, which would have provided immense context to the “outer” -omics upon which our analysis focused, and future analysis incorporating host genome data would add to these results by allowing comparison of the outer -omics to standard polygenic risk scores for IBD. Additionally, no information was available regarding the existence of any relatives within the dataset, which could have confounded results, as our analyses assumed all participants were not related or living in the same location.

## Conclusions

Not only did we successfully implement a poly-omic risk score framework across four -omics layers, each of our individual models identified features known to associate with IBD risk while providing new insights into features that may influence or be influenced by IBD. The majority of IBD models do not utilize multiple -omics layers, focusing instead on host genetics or solely microbiome taxonomic composition.^56^ Those that utilize other information often focus on predicting flare-up and other clinical outcomes among individuals with IBD.^39^ This study is unique in that it combines -omics layers to predict the presence of disease.

Our results provide a framework for an interpretable comparison of single-omic models in multi-omic contexts, with particular relevance to the gut microbiome and complex phenotypes. Our work suggests that some single-omic models (in this case MBL and VRM) are more predictive than others, though this is likely phenotype-dependent. Each -omic model provides a combination of unique and redundant information, relative to other -omic models, and a combination of single-omic models may often yield improvements in predictive accuracy. Such methods are extendable and customizable to the context of interest through the use of non-linear methods, alternative data transformations, and interactions between features. There are generally tradeoffs between model simplicity, predictive accuracy, and interpretability, and future studies should carefully consider the primary goals of their model. Multi-omic modeling approaches that connect the dots along the continuum from genes to their environment will be paramount to identifying novel insights for health and disease.

## Acknowledgements

We would like to thank Kristin Powell and Stephanie Hoyt for their support for this project, as well as Tom Cech and the BioFrontiers Institute for their support of the Interdisciplinary Quantitative Biology program. We’d also like to thank Lukas Buecherl and Casey Martin for providing manuscript feedback and the Human Microbiome Project 2 team for data collection, processing, and public access and HMP2 participants.

## Author Order

CA and JS are co-first authors. Co-first author order was determined by coin flip.

