## Supplemental Material for "Poly-omic risk scores predict inflammatory bowel disease diagnosis"

### Supplementary Information

**Supplementary Table 1. Dataset size and split for training and validation.** The table shows the number of samples belonging to each respective category, with the number of participants those samples correspond to shown in parentheses. One participant was removed during the split to training and validation datasets due to missing data. *Because the multi-omic model needed all four -omic data types, only samples that included all four -omic data types could be retained. As a result, samples missing one or more -omics were discarded, and validation samples retained for multi-omic testing are shown in the far right column. Abbreviations: QC, quality control; LASSO, least absolute shrinkage and selection operator; MGN, metagenomics; MTS, metatranscriptomics; VRM, viromics; MBL, metabolomics.

| Dataset | N features passing QC | N features selected by LASSO | Total samples (participants) | Training samples (participants) | Validation samples (participants) | Validation samples (participants) retained for multi-omic testing* |
|---|---|---|---|---|---|---|
| Metagenomics (MGN) | 237 | 14 | 1627 (130) | 1210 (100) | 417 (30) | 114 (30) |
| Metatranscriptomics (MTS) | 280 | 23 | 804 (109) | 587 (79) | 217 (30) | 114 (30) |
| Metabolomics (MBL) | 596 | 14 | 546 (130) | 374 (76) | 172 (30) | 114 (30) |
| Viromics (VRM) | 9 | 6 | 703 (105) | 493 (75) | 210 (30) | 114 (30) |

**Supplementary Table 2**. **Measures of accuracy for different modeling approaches.** A general improvement in accuracy (larger AUCs and ORs) is observed moving down the table as model progression moves from the baseline model (covariates only), to a model with only the standardized score, to a model that includes the standardized score + age + sex. Abbreviations: MGN, metagenomics; MTS, metatranscriptomics; VRM, viromics; MBL, metabolomics; AUC, area under the receiver operating characteristic curve; OR, odds ratio; CI, confidence interval.

| | **MGN** | **MTS** | **MBL** | **VRM** | **MGN + MTS + MBL + VRM** |
|---|---|---|---|---|---|
| Baseline Model AUC [95% CI], Nagelkerke's $R^2$ | 0.43 [0.19, 0.67], 0.08 | 0.38 [0.15, 0.62], 0.04 | 0.47 [0.25, 0.69], 0.01 | 0.47 [0.23, 0.71], 0.02 | NA |
| IBD ~ score | 0.70 [0.50, | 0.64 [0.42, | 0.78 [0.59, | 0.80 [0.63, | 0.77 [0.58, |

| | | | | | |
|---|---|---|---|---|---|
| AUC [95% CI], Nagelkerke's $R^2$ | 0.91], 0.04 | 0.86], 0.12 | 0.98], 0.32 | 0.96], 0.31 | 0.96], 0.37 |
| IBD ~ score OR [95% CI], p-value | 1.45 [0.63, 3.37], 0.4 | 3.12 [0.72, 13.53], 0.1 | 4.21 [1.34, 13.19], 0.01 | 5.94 [1.38, 25.55], 0.02 | 3.13 [1.19, 8.26], 0.02 |
| IBD ~ score + age + sex AUC [95% CI], Nagelkerke's R2 | 0.66 [0.44, 0.87], 0.12 | 0.73 [0.53, 0.92], 0.20 | 0.82 [0.66, 0.98], 0.40 | 0.83 [0.68, 0.98], 0.43 | 0.80 [0.63, 0.98], 0.46 |
| IBD ~ score + age + sex OR [95% CI], p-value | 1.24 [0.50, 3.09], 0.6 | 2.89 [0.62, 13.52], 0.2 | 4.30 [1.32, 13.98], 0.02 | 9.28 [1.53, 56.13], 0.02 | 3.14 [1.10, 9.00], 0.03 |

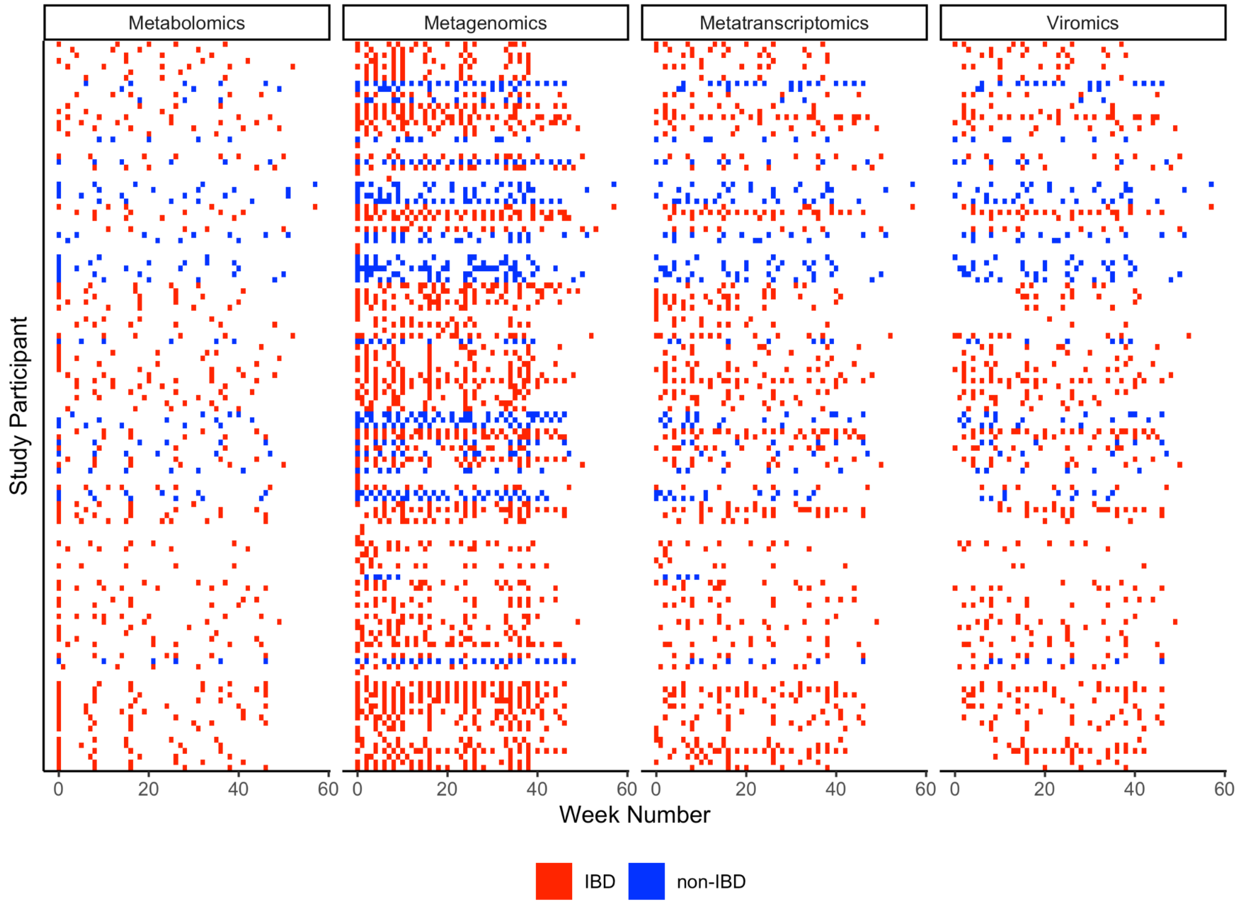


**Supplementary Figure 1.** The longitudinal sampling frequency of each subject over a 57 week time period for the 4 analyzed datasets. Metabolomics had an average of 10.3 weeks between samples, metagenomics an average of 3.3 weeks, metatranscriptomics an average of 6.5 weeks, and viromics an average of 7.4 weeks. Abbreviations: IBD, inflammatory bowel disease.

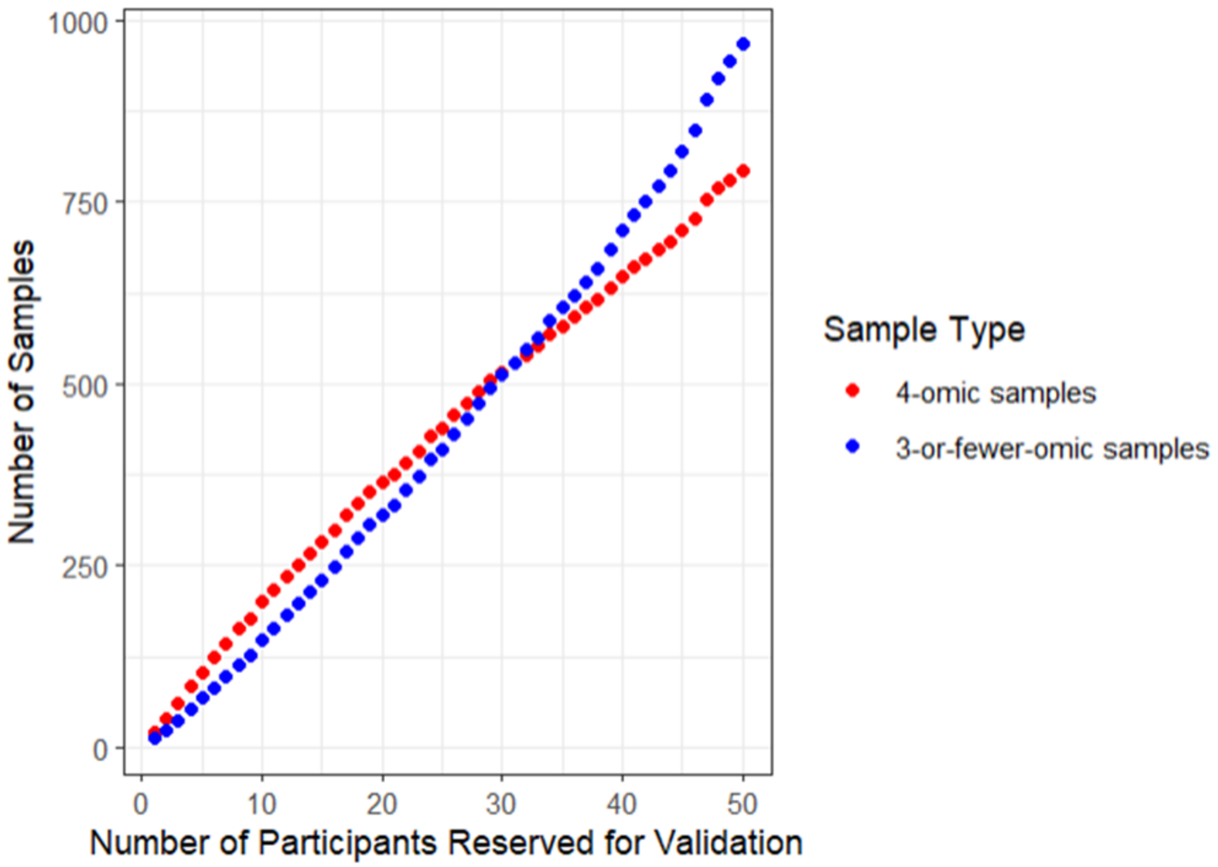


**Supplementary Figure 2. 4-omic samples and 3-or-fewer-omic samples when splitting the training and testing datasets.** The x-axis shows the number of participants chosen for testing (chosen in order of participants with the most 4-omics samples). The y-axis shows the cumulative number of samples from those participants. Red dots represent the number of samples containing all four -omics, whereas blue dots represent the number of samples that do not contain all four -omics of interest. We chose to reserve the 30 participants (23% of individuals) with the most samples containing all four -omics to maximize the number of usable multi-omic testing samples, as samples without all four -omics could not be used in the combined multi-omic model.

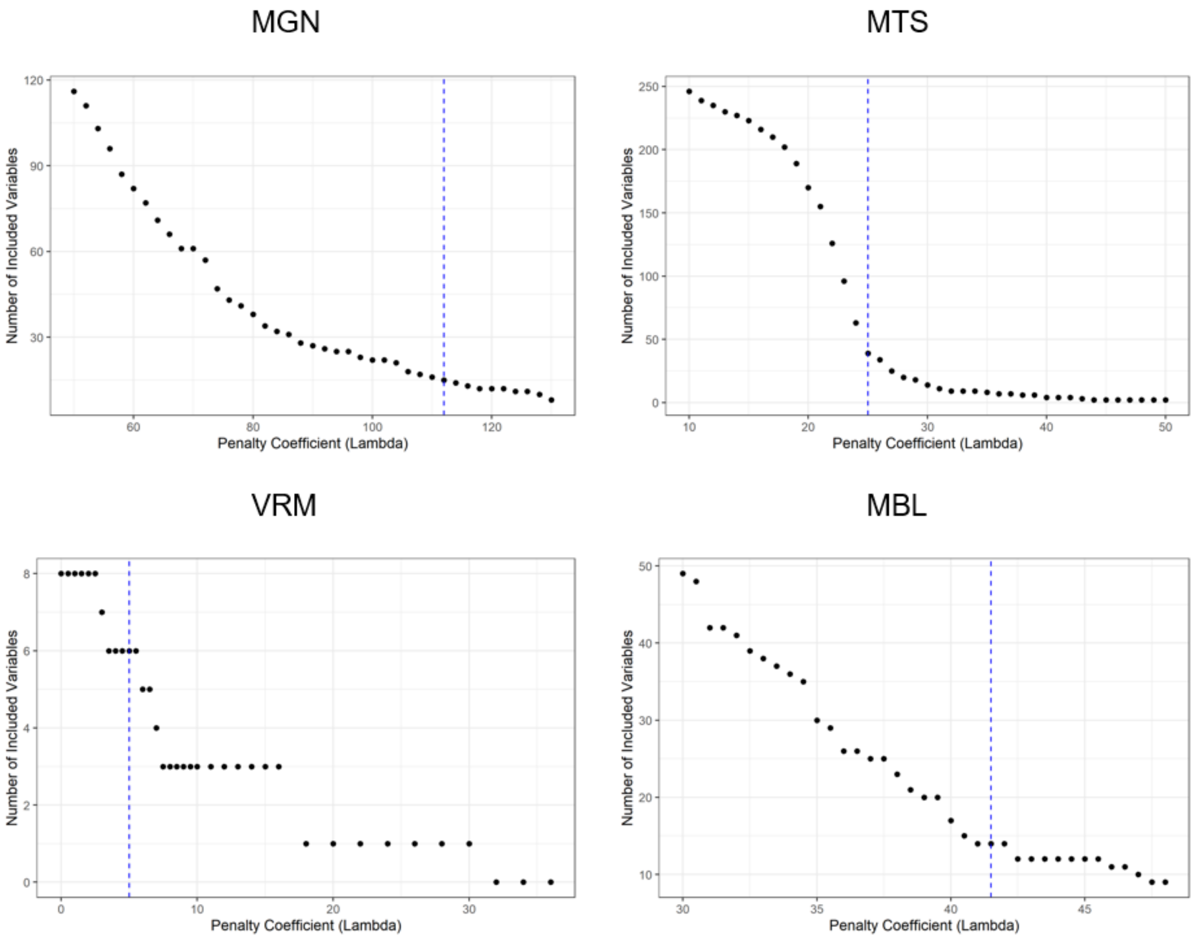


**Supplementary Figure 3. Variables retained by mixed effect LASSO using varying penalty parameters.** Increasing values along the x-axis represent increased penalty for variable inclusion in the final model, and the y-axis represents the number of features retained for a given penalty coefficient. A vertical dashed line was drawn at the chosen “elbow” for each -omics data type to describe the lambda that was used going forward. The number of variables shown along the y-axis only accounts for fixed effects and does not include the participant metadata covariates, such as age, sex, antibiotic use, or race. Abbreviations: MGN, metagenomics; MTS, metatranscriptomics; VRM, viromics; MBL, metabolomics; LASSO, least absolute shrinkage and selection operator.

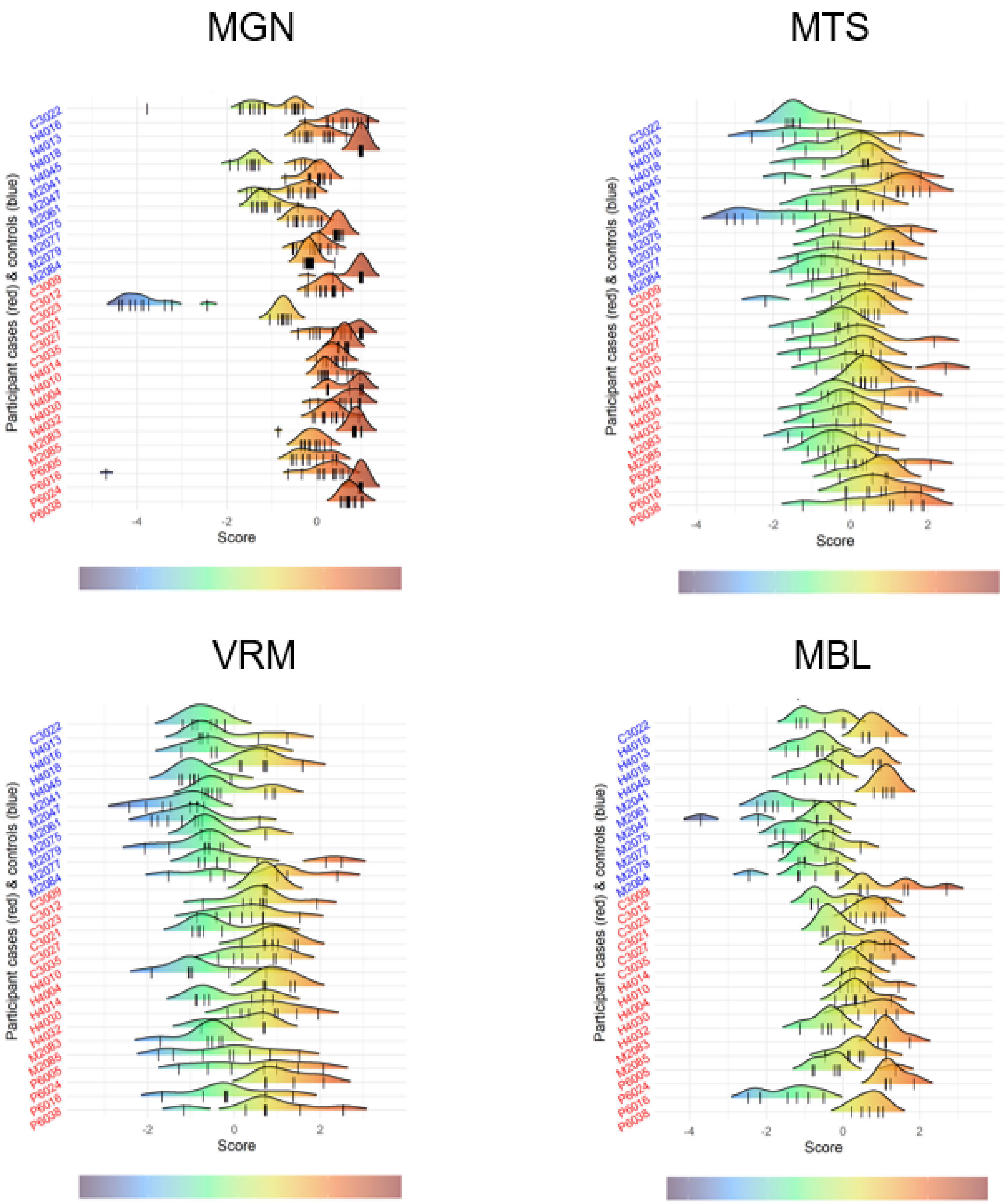


**Supplementary Figure 4. Standardized scores per participant derived for each of the samples reserved for model testing.** The score distribution across each individual's samples is depicted on the x-axis, where controls are labeled in blue and cases are labeled in red. The scores (represented as one black tick mark per sample) were averaged for each individual before assessing model accuracy or combining with scores across other -omics. MTS tends to have the most variability among scores within individuals whereas MGN tends to have the least variability among scores within individuals. This distribution plot helps illustrate how dynamic the

-omics layers are between sampling timepoints. Abbreviations: MGN, metagenomics; MTS, metatranscriptomics; VRM, viromics; MBL, metabolomics.

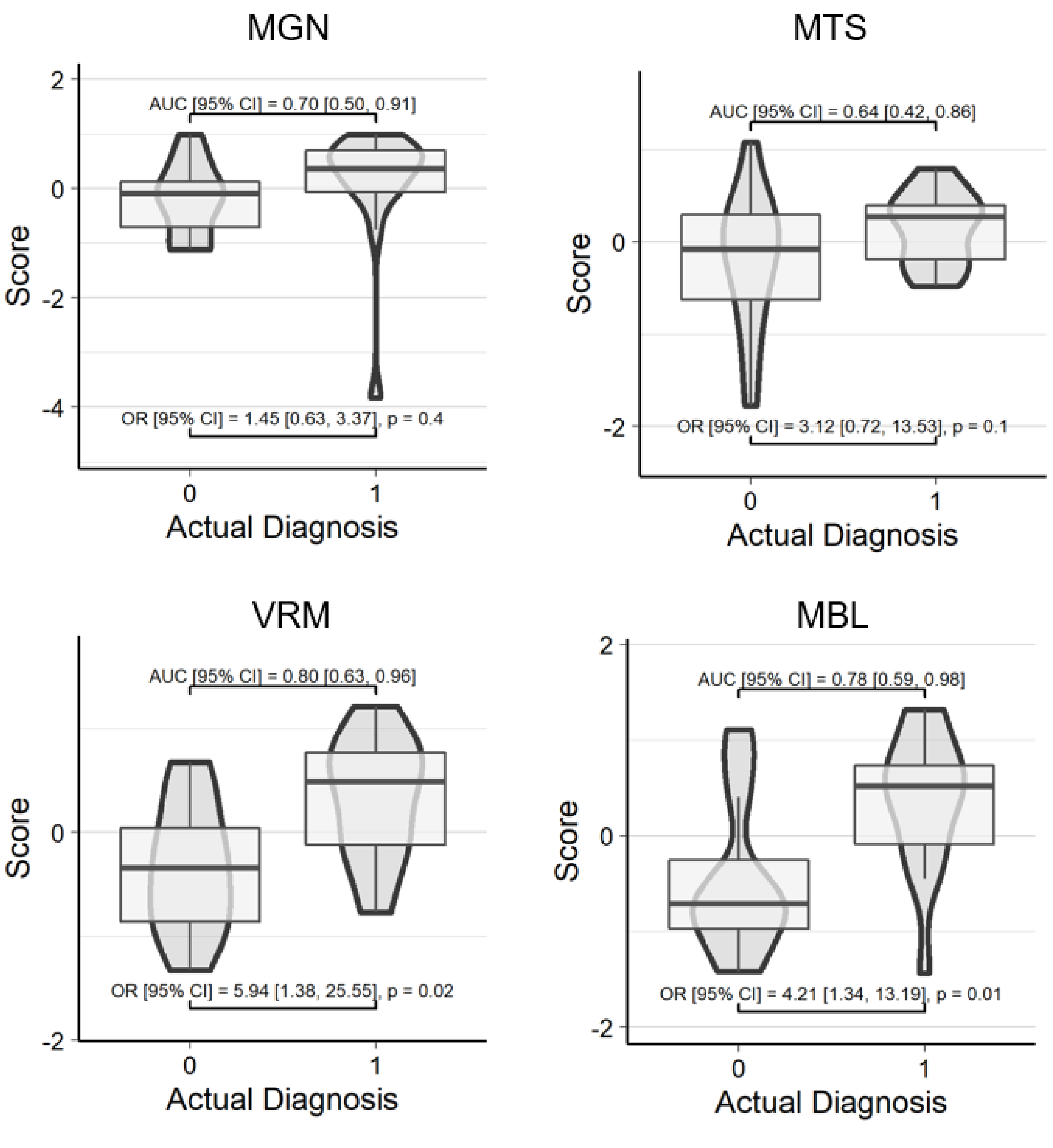


**Supplementary Figure 5. Risk scores without covariates predict IBD diagnosis.** Z-score transformed risk scores (averaged for each participant) on the y-axis are plotted against actual diagnosis on the x-axis for the validation dataset. In contrast to Figure 1, AUC and OR were calculated with score as the only predictor (diagnosis ~ score). Each of the four scores shown were calculated using feature weights from a LASSO-identified mixed effect logistic regression trained in a separate set of samples/individuals: diagnosis ~ features + age + sex + race + antibiotic use + (1|site) + (1|participant ID). An actual diagnosis value of 1 indicates presence of

IBD. Abbreviations: MGN, metagenomics; MTS, metatranscriptomics; VRM, viromics; MBL, metabolomics; AUC, area under the receiver operating characteristic curve; OR, odds ratio; CI, confidence interval; IBD, inflammatory bowel disease.

**Supplementary Figures 6-19.** Spaghetti plots (panel A) depicting the transformed relative abundance of each LASSO-selected metagenomics species longitudinally for the individuals in the testing dataset. Smoothed conditional means are shown with shaded 95% confidence intervals colored blue for controls and red for cases to help view the overall patterns between groups. Panel B shows the participant mean transformed relative abundance by case/control status. In both panel A and panel B participants' data are colored according to the legend. Abbreviations: LASSO, least absolute shrinkage and selection operator.

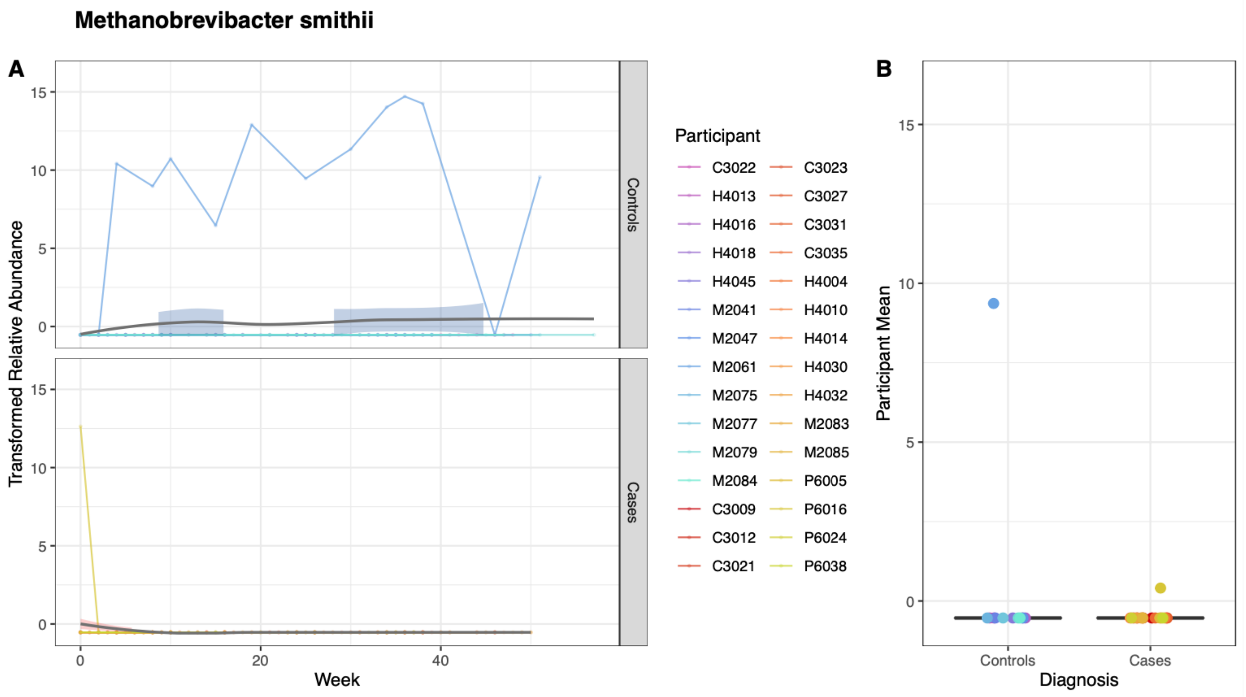


**Supplementary Figure 6.** Figure legend above.

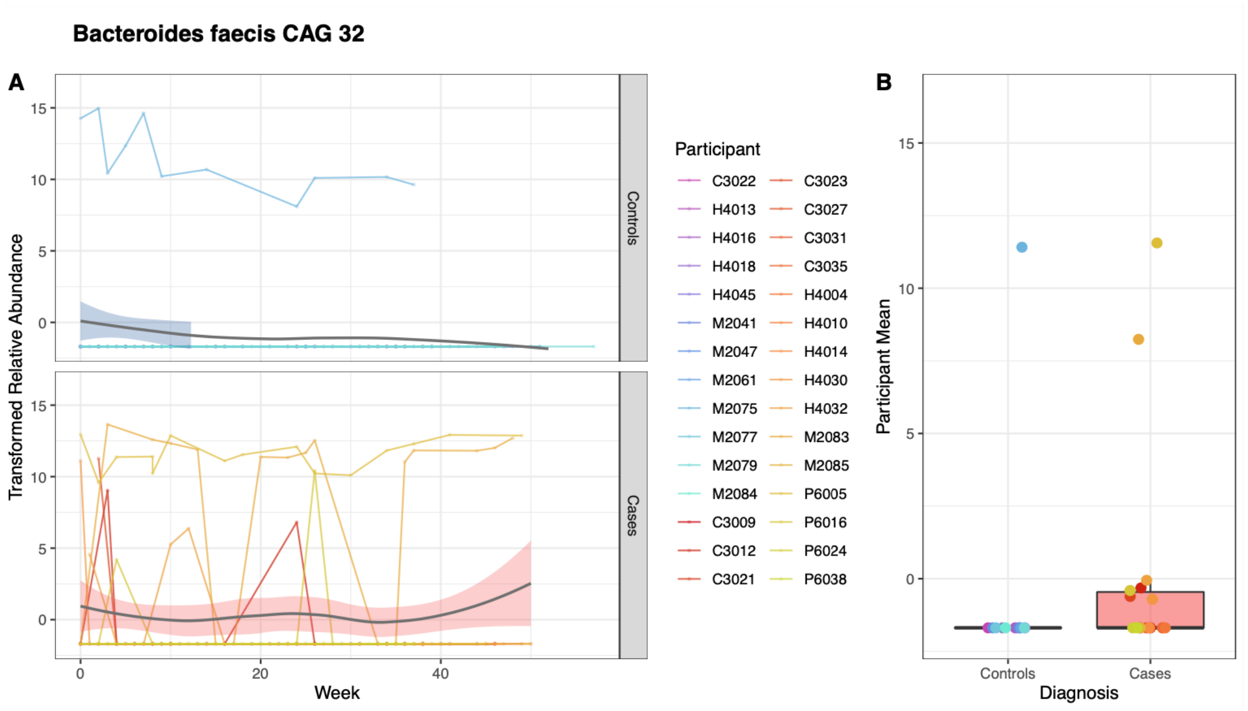


**Supplementary Figure 7.** Figure legend above.

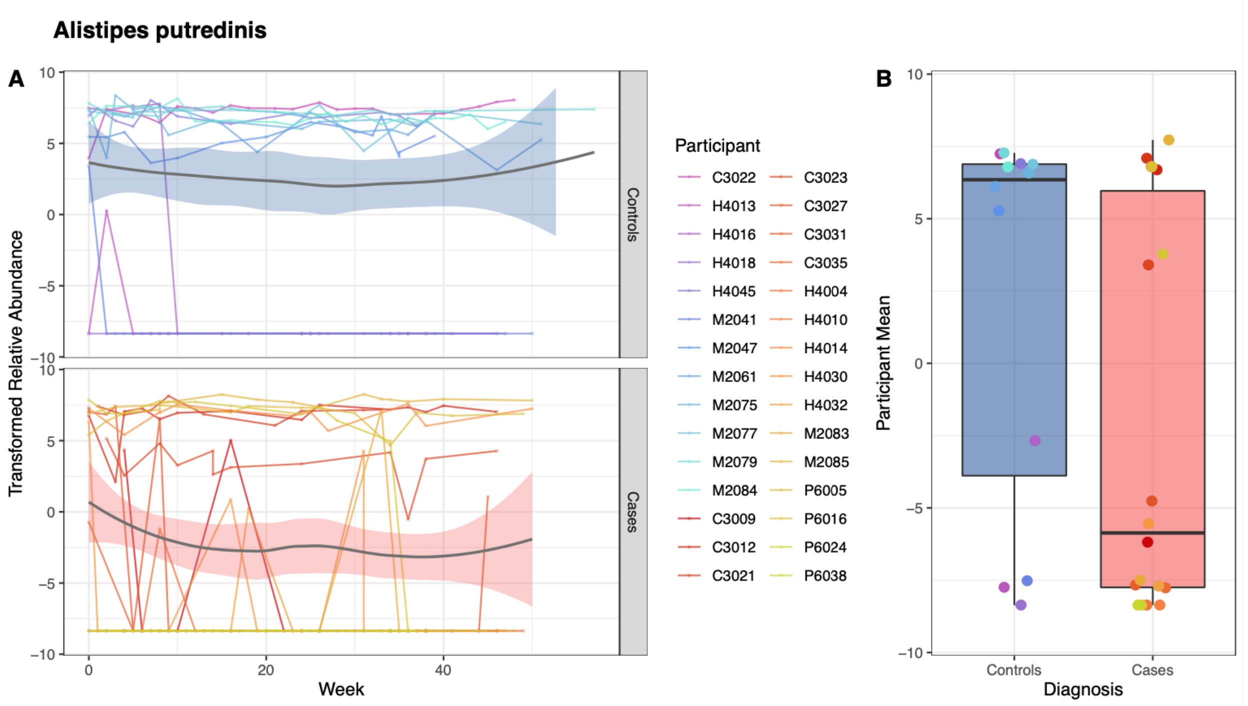


**Supplementary Figure 8.** Figure legend above.

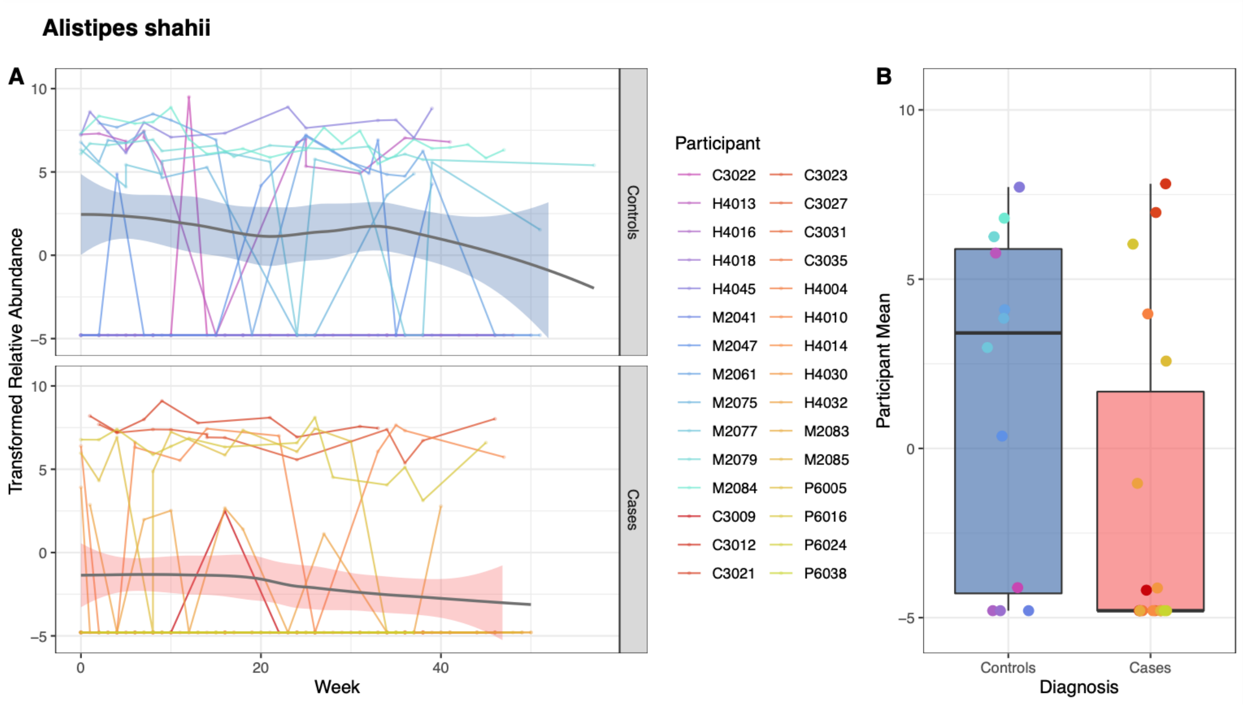


**Supplementary Figure 9.** Figure legend above.

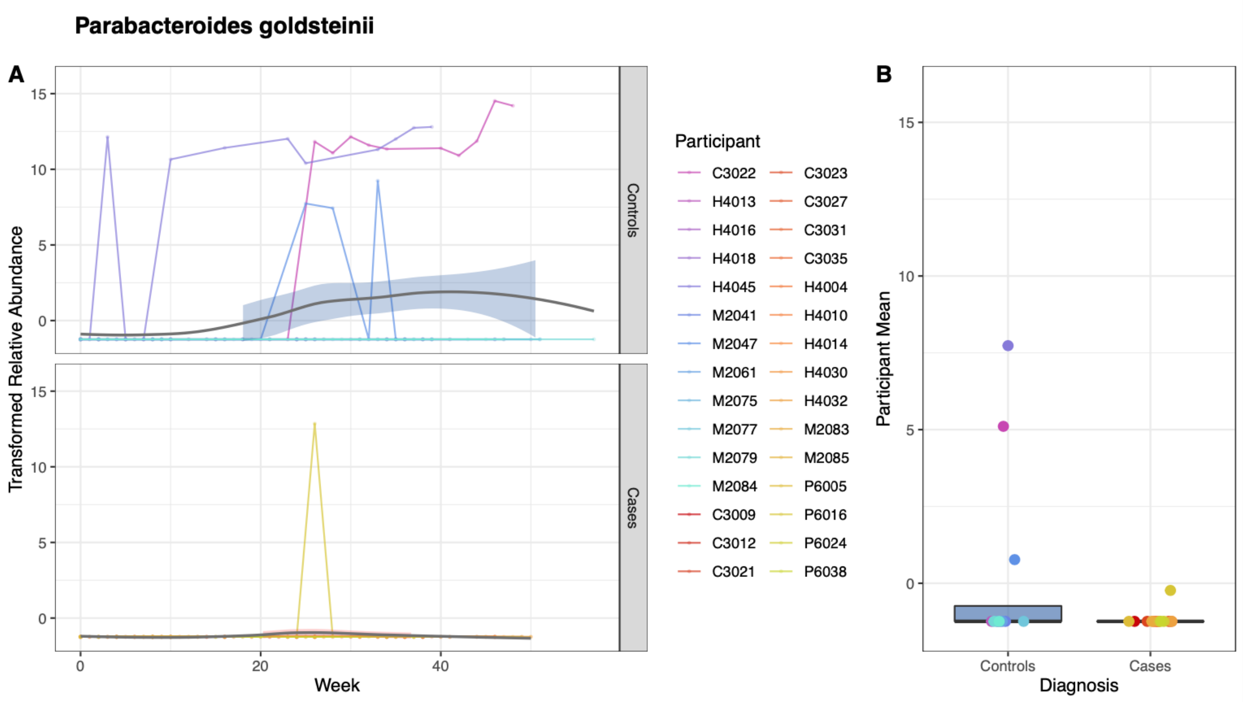


**Supplementary Figure 10.** Figure legend above.

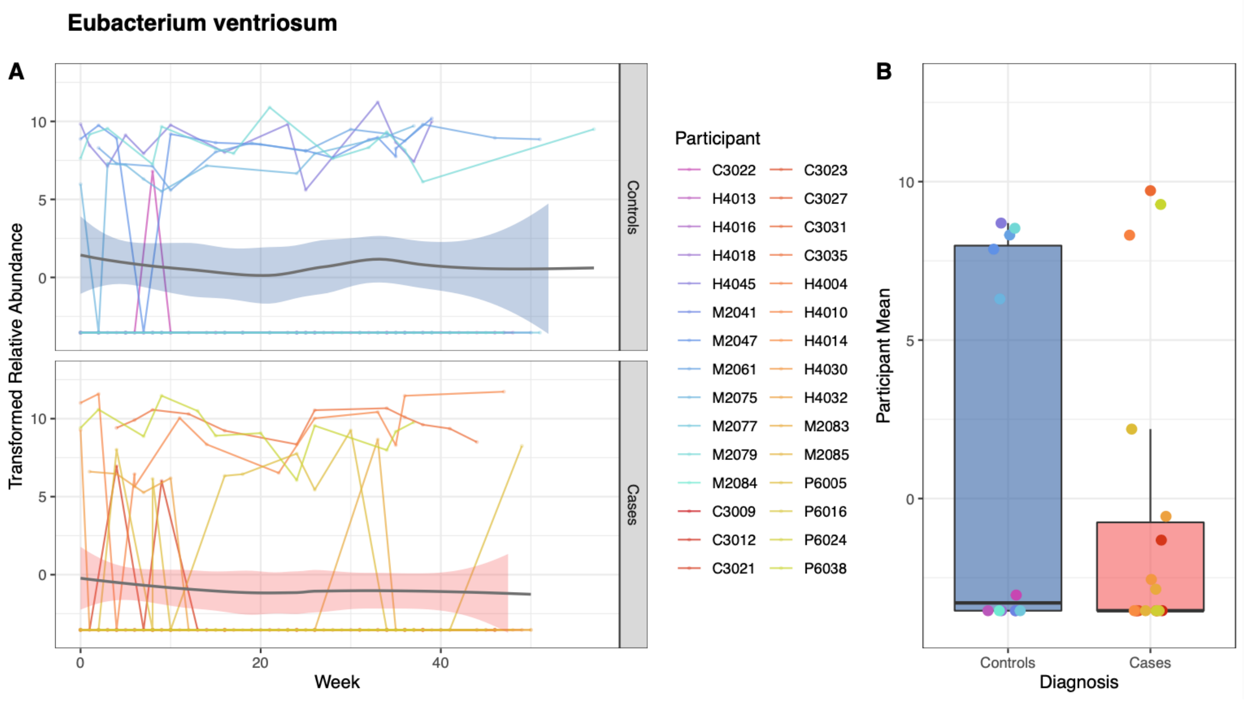


**Supplementary Figure 11.** Figure legend above.

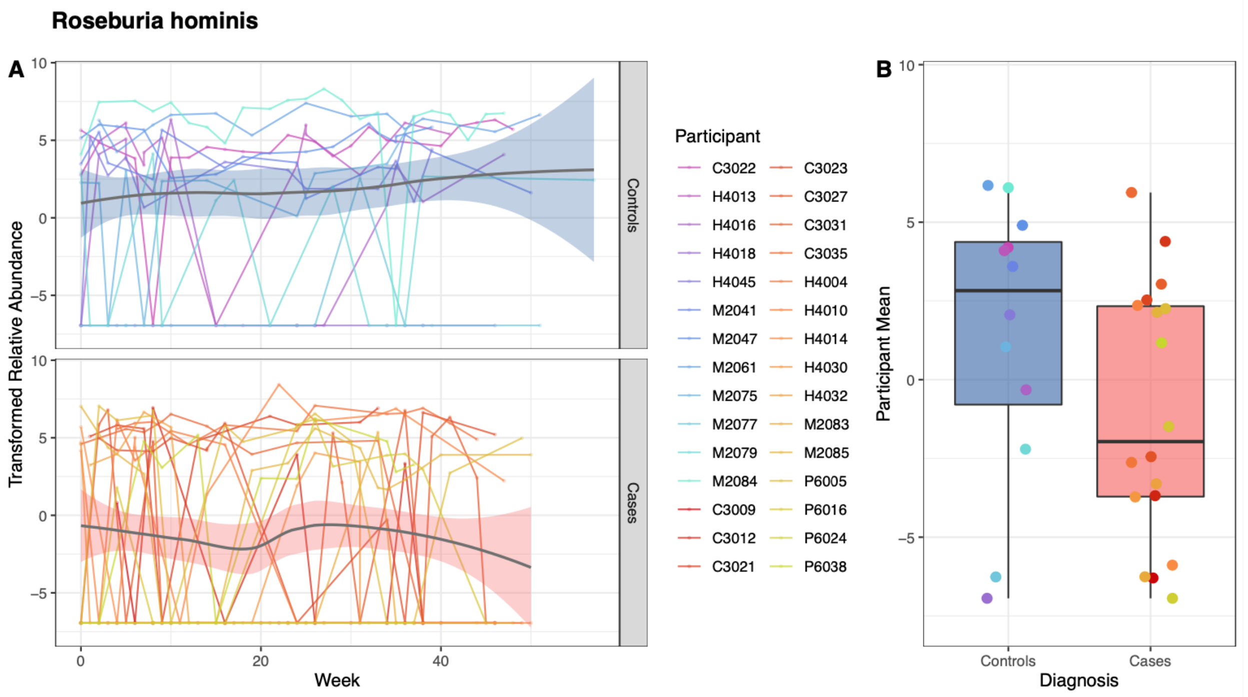


**Supplementary Figure 12.** Figure legend above.

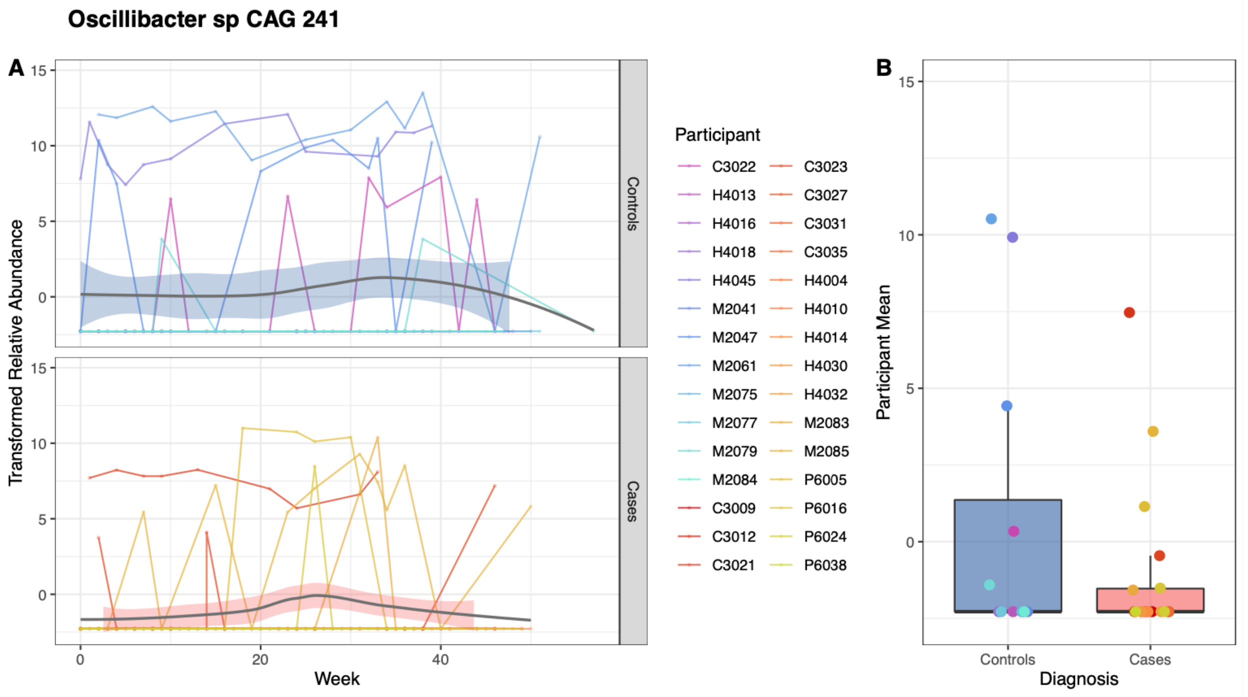


**Supplementary Figure 13.** Figure legend above.

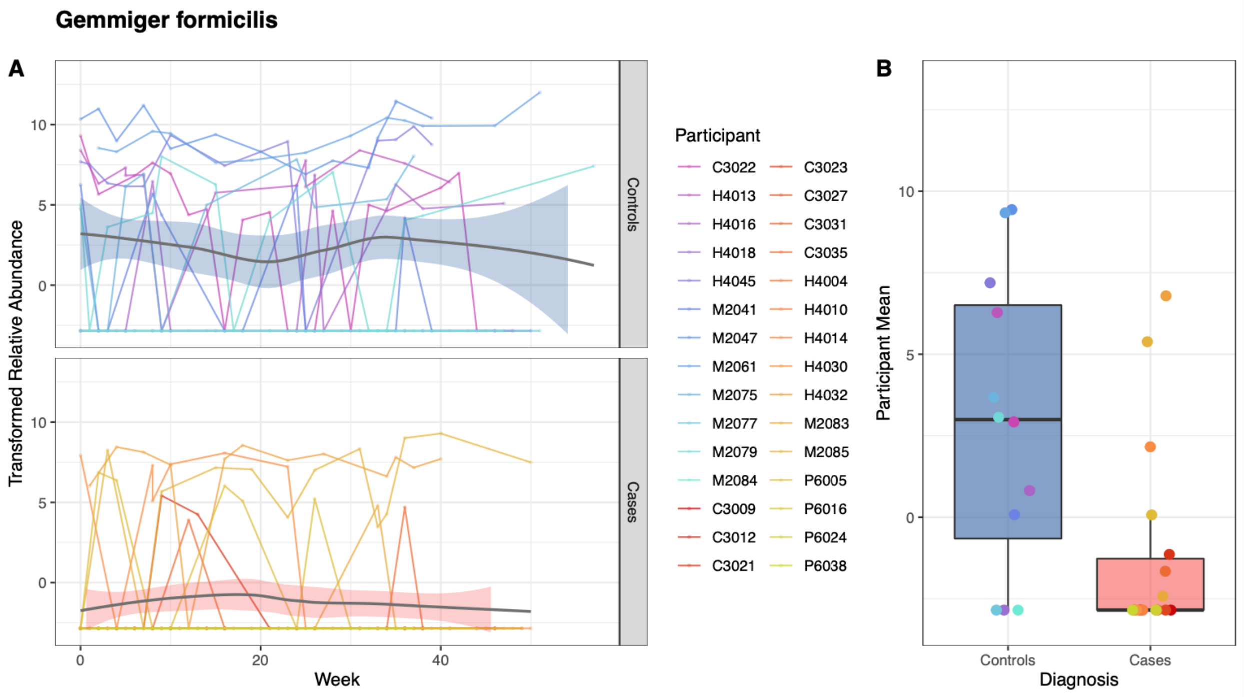


**Supplementary Figure 14.** Figure legend above.

Eubacterium siraeum

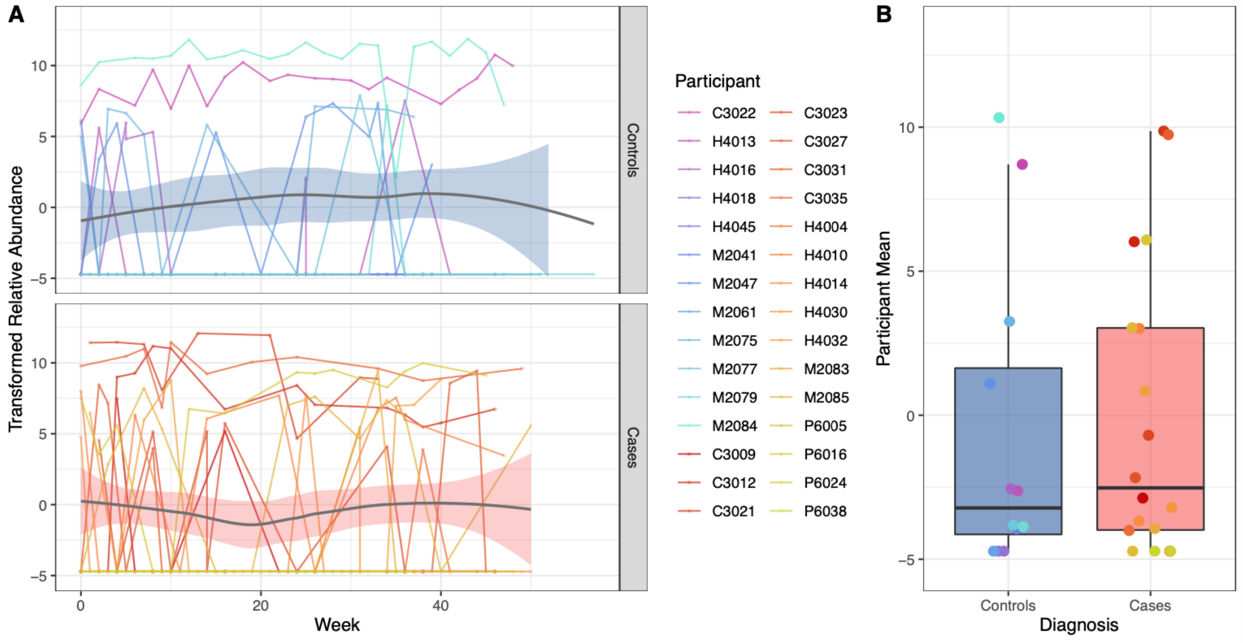


**Supplementary Figure 15.** Figure legend above.

Ruminococcus bromii

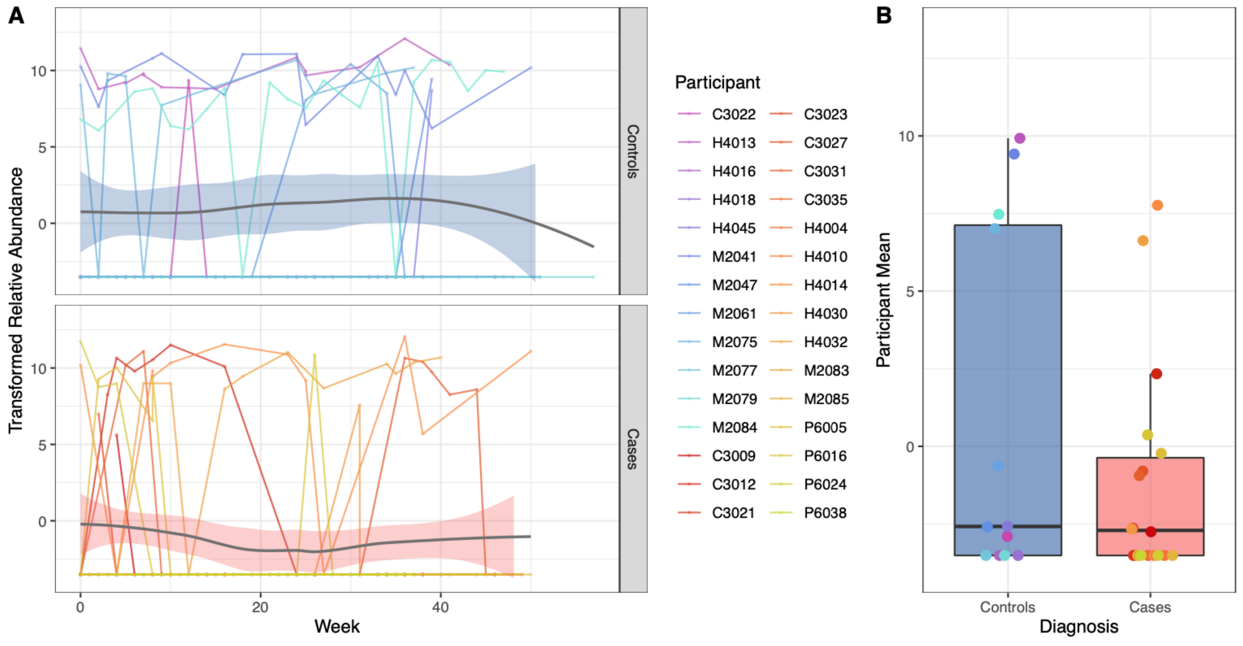


**Supplementary Figure 16.** Figure legend above.

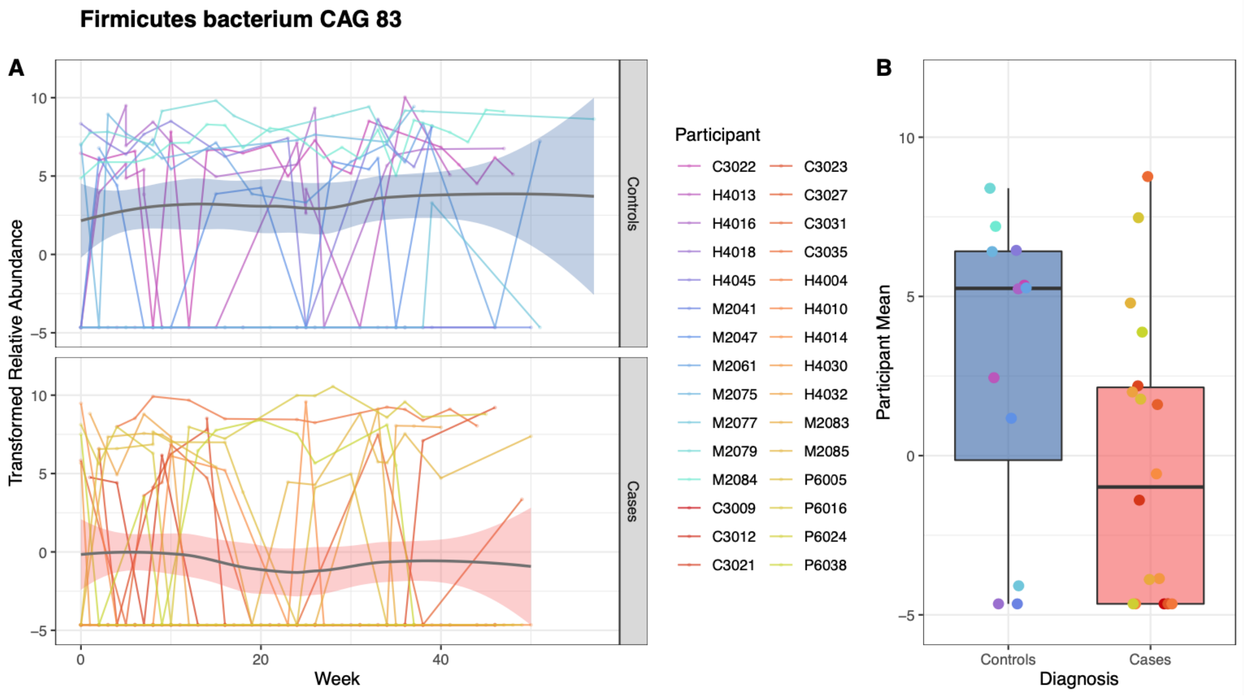


**Supplementary Figure 17.** Figure legend above.

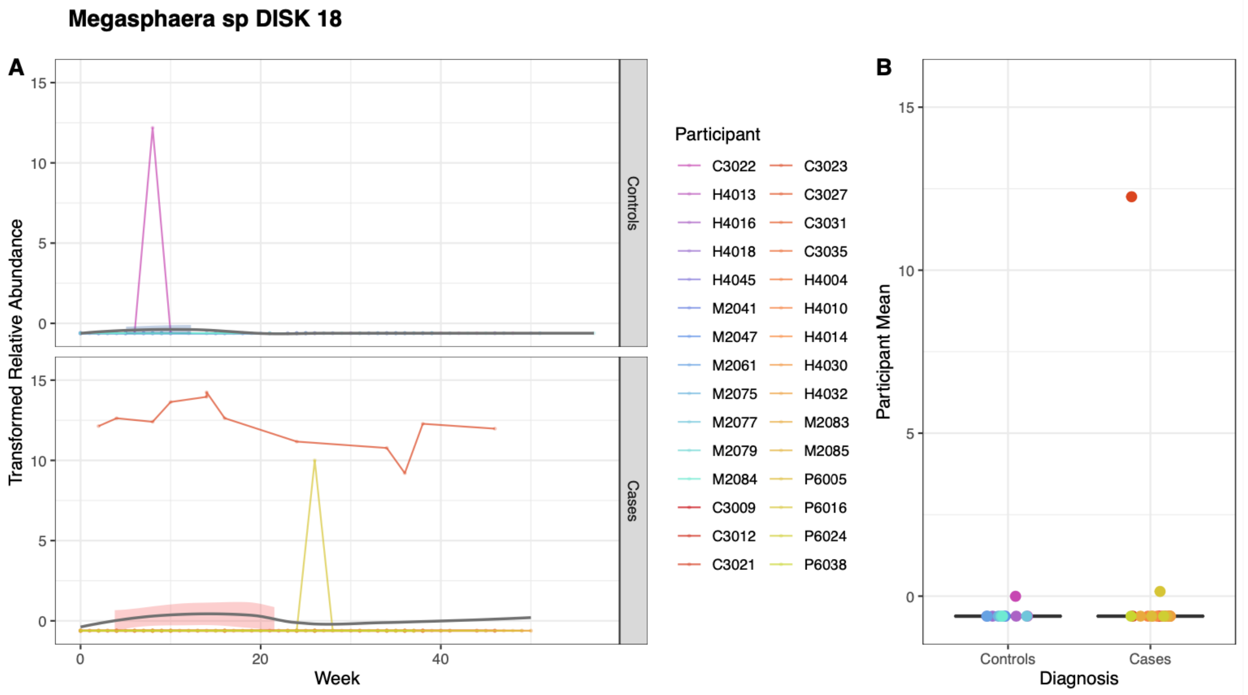


**Supplementary Figure 18.** Figure legend above.

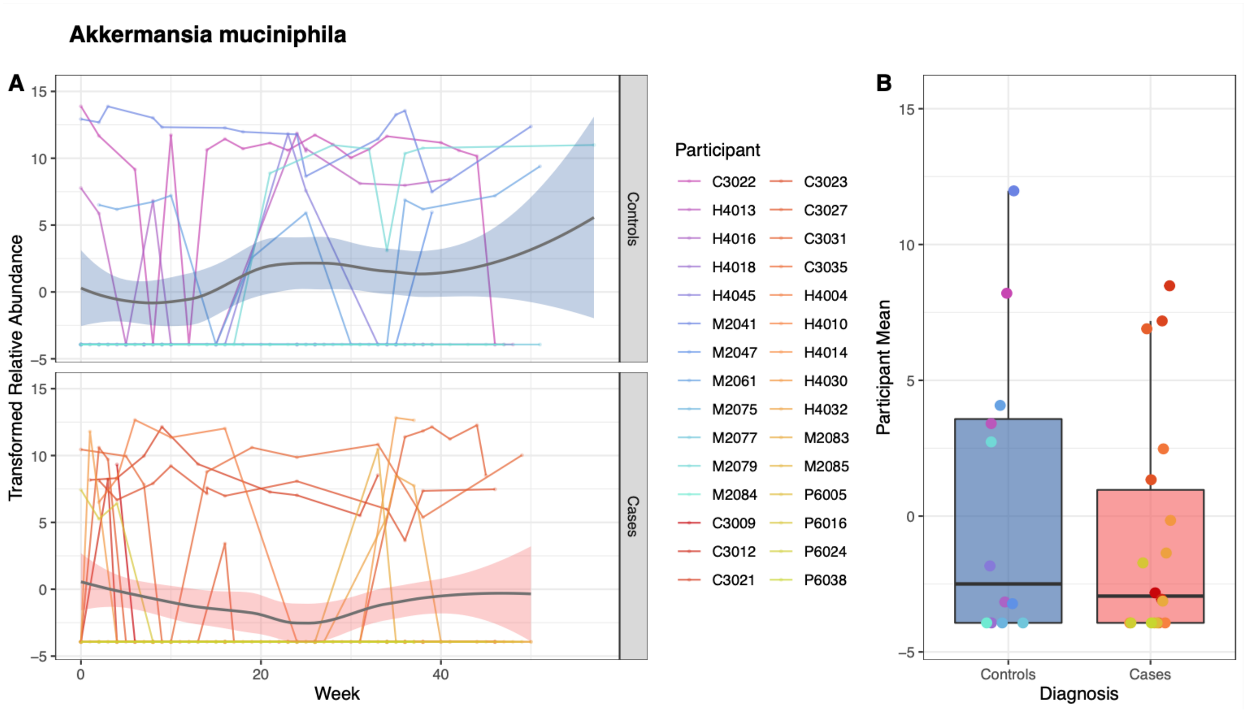


**Supplementary Figure 19.** Figure legend above.

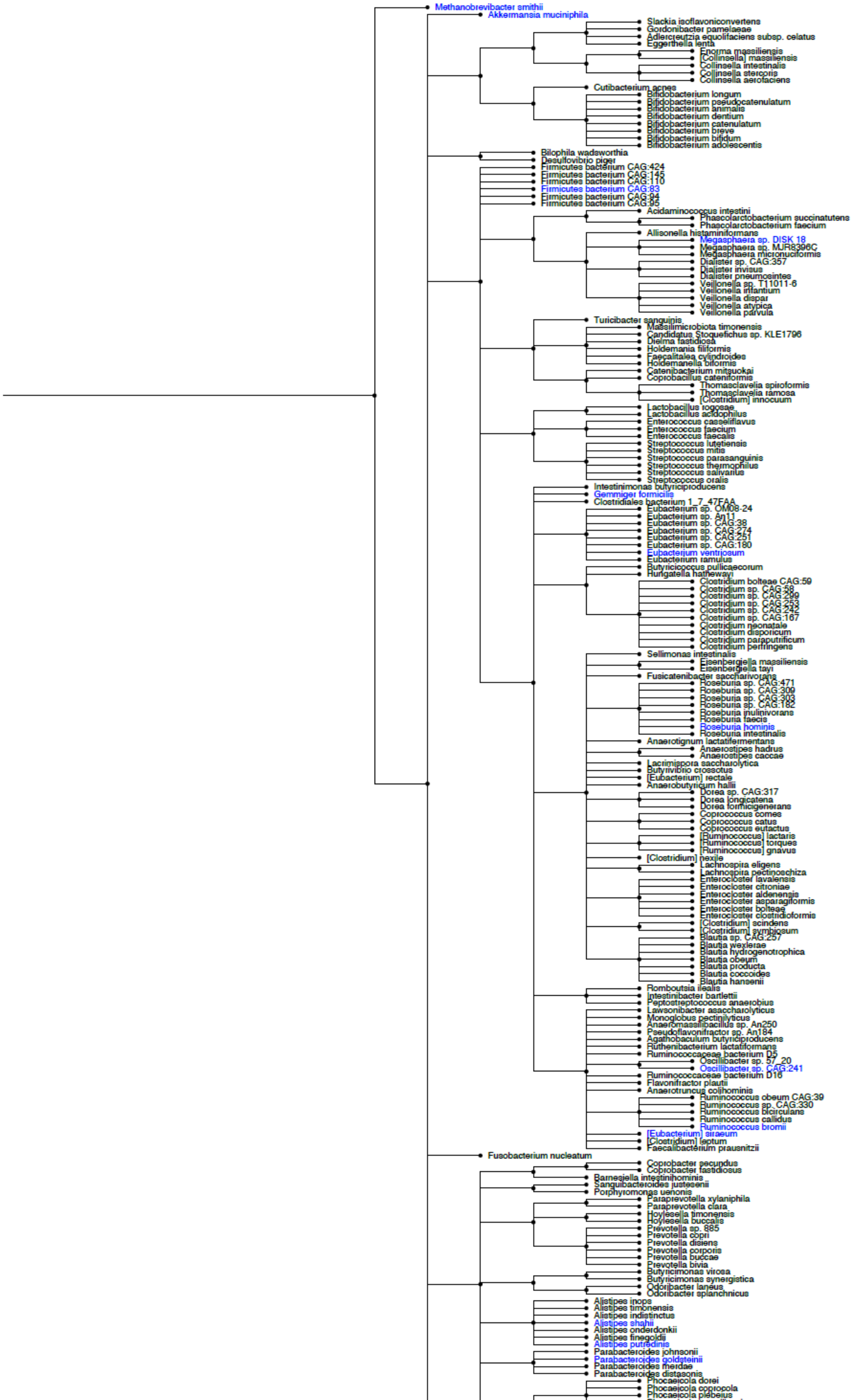
Methanobrevibacter smithii
Akkermansia muciniphila
Slackia isoflavoniconvertens
Gordonibacter pamelaeae
Adlercreutzia equolifaciens subsp. celatus
Eggerthella lenta
Enorma massiliensis
[Collinsella] massiliensis
Collinsella intestinalis
Collinsella stercoris
Collinsella aerofaciens
Cutibacterium acnes
Bifidobacterium longum
Bifidobacterium pseudocatenulatum
Bifidobacterium animalis
Bifidobacterium dentium
Bifidobacterium catenulatum
Bifidobacterium breve
Bifidobacterium bifidum
Bifidobacterium adolescentis
Bilophila wadsworthia
Desulfovibrio piger
Firmicutes bacterium CAG:424
Firmicutes bacterium CAG:145
Firmicutes bacterium CAG:170
Firmicutes bacterium CAG:83
Firmicutes bacterium CAG:94
Firmicutes bacterium CAG:95
Acidaminococcus intestini
Phascolarctobacterium succinatutens
Phascolarctobacterium faecium
Allisonella histaminiformans
Megasphaera sp. DISK 18
Megasphaera sp. MJR8396C
Megasphaera micronuciformis
Dialister sp. CAG:357
Dialister invisus
Dialister pneumosintes
Veillonella sp. T11011-6
Veillonella infantium
Veillonella dispar
Veillonella atypica
Veillonella parvula
Turicibacter sanguinis
Massilimicrobiota timonensis
Candidatus Stoquefichus sp. KLE1796
Dielma fastidiosa
Holdemania filiformis
Faecalitalea cylindroides
Holdemanella biformis
Catenibacterium mitsuokai
Coprobacillus cateniformis
Thomasclavelia spiroformis
Thomasclavelia ramosa
[Clostridium] innocuum
Lactobacillus rogosae
Lactobacillus acidophilus
Enterococcus casseliflavus
Enterococcus faecium
Enterococcus faecalis
Streptococcus lutetiensis
Streptococcus mitis
Streptococcus parasanguinis
Streptococcus thermophilus
Streptococcus salivarius
Streptococcus oralis
Intestinimonas butyriciproducens
Gemmiger formicilis
Clostridiales bacterium 1_7_47FAA
Eubacterium sp. OM08-24
Eubacterium sp. An11
Eubacterium sp. CAG:38
Eubacterium sp. CAG:274
Eubacterium sp. CAG:251
Eubacterium sp. CAG:180
Eubacterium ventriosum
Eubacterium ramulus
Butyricicoccus pullicaecorum
Hungatella hathewayi
Clostridium bolteae CAG:59
Clostridium sp. CAG:58
Clostridium sp. CAG:299
Clostridium sp. CAG:253
Clostridium sp. CAG:242
Clostridium sp. CAG:167
Clostridium neonatale
Clostridium disporicum
Clostridium paraputrificum
Clostridium perfringens
Sellimonas intestinalis
Eisenbergiella massiliensis
Eisenbergiella tayi
Fusicatenibacter saccharivorans
Roseburia sp. CAG:471
Roseburia sp. CAG:309
Roseburia sp. CAG:303
Roseburia sp. CAG:182
Roseburia inulinivorans
Roseburia faecis
Roseburia hominis
Roseburia intestinalis
Anaerotignum lactatifermentans
Anaerostipes hadrus
Anaerostipes caccae
Lacrimispora saccharolytica
Butyrivibrio crossotus
[Eubacterium] rectale
Anaerobutyricum hallii
Dorea sp. CAG:317
Dorea longicatena
Dorea formicigenerans
Coprococcus comes
Coprococcus catus
Coprococcus eutactus
[Ruminococcus] lactaris
[Ruminococcus] torques
[Ruminococcus] gnavus
[Clostridium] nexile
Lachnospira eligens
Lachnospira pectinoschiza
Enterocloster lavalensis
Enterocloster citroniae
Enterocloster aldenensis
Enterocloster asparagiformis
Enterocloster bolteae
Enterocloster clostridioformis
[Clostridium] scindens
[Clostridium] symbiosum
Blautia sp. CAG:257
Blautia wexlerae
Blautia hydrogenotrophica
Blautia obeum
Blautia producta
Blautia coccoides
Blautia hansenii
Romboutsia ilealis
Intestinibacter bartlettii
Peptostreptococcus anaerobius
Lawsonibacter asaccharolyticus
Monoglobus pectinilyticus
Anaeromassilibacillus sp. An250
Pseudoflavonifractor sp. An184
Agathobaculum butyriciproducens
Ruthenibacterium lactatiformans
Ruminococcaceae bacterium D5
Oscillibacter sp. 57_20
Oscillibacter sp. CAG:241
Ruminococcaceae bacterium D16
Flavonifractor plautii
Anaerotruncus colihominis
Ruminococcus obeum CAG:39
Ruminococcus sp. CAG:330
Ruminococcus bicirculans
Ruminococcus callidus
Ruminococcus bromii
[Eubacterium] siraeum
[Clostridium] leptum
Faecalibacterium prausnitzii
Fusobacterium nucleatum
Coprobacter secundus
Coprobacter fastidiosus
Barnesiella intestinihominis
Sanguibacteroides justesenii
Porphyromonas uenonis
Paraprevotella xylaniphila
Paraprevotella clara
Hoylesella timonensis
Hoylesella buccalis
Prevotella sp. 885
Prevotella copri
Prevotella disiens
Prevotella corporis
Prevotella buccae
Prevotella bivia
Butyricimonas virosa
Butyricimonas synergistica
Odoribacter laneus
Odoribacter splanchnicus
Alistipes inops
Alistipes timonensis
Alistipes indistinctus
Alistipes shahii
Alistipes onderdonkii
Alistipes finegoldii
Alistipes putredinis
Parabacteroides johnsonii
Parabacteroides goldsteinii
Parabacteroides merdae
Parabacteroides distasonis
Phocaeicola dorei
Phocaeicola coprocola
Phocaeicola plebeius

**Supplementary Figure 20:** A phylogenetic tree illustrating the hierarchical taxonomical relationships for bacteria species that were present in the MGN dataset and the 14 LASSO-selected features colored in blue. The phylogeny was constructed using the NCBI Common Tree database using FigTree v1.4.4. 217 out of 237 bacteria species in the MGN dataset matched the reference NCBI Common Tree database and 20 were included in the phylogeny as heterotypic synonym. Among the 14 LASSO-selected features that were used in the MGN model, we observed that *Alistipes putredinis* and *Alistipes shahii* are the most closely related species and *Methanobrevibacter smithii* appeared to be more distantly related to the other 13 features in the phylogenetic tree. Abbreviations: MGN, metagenomics; NCBI, National Center of Biotechnology Information; LASSO, least absolute shrinkage and selection operator.

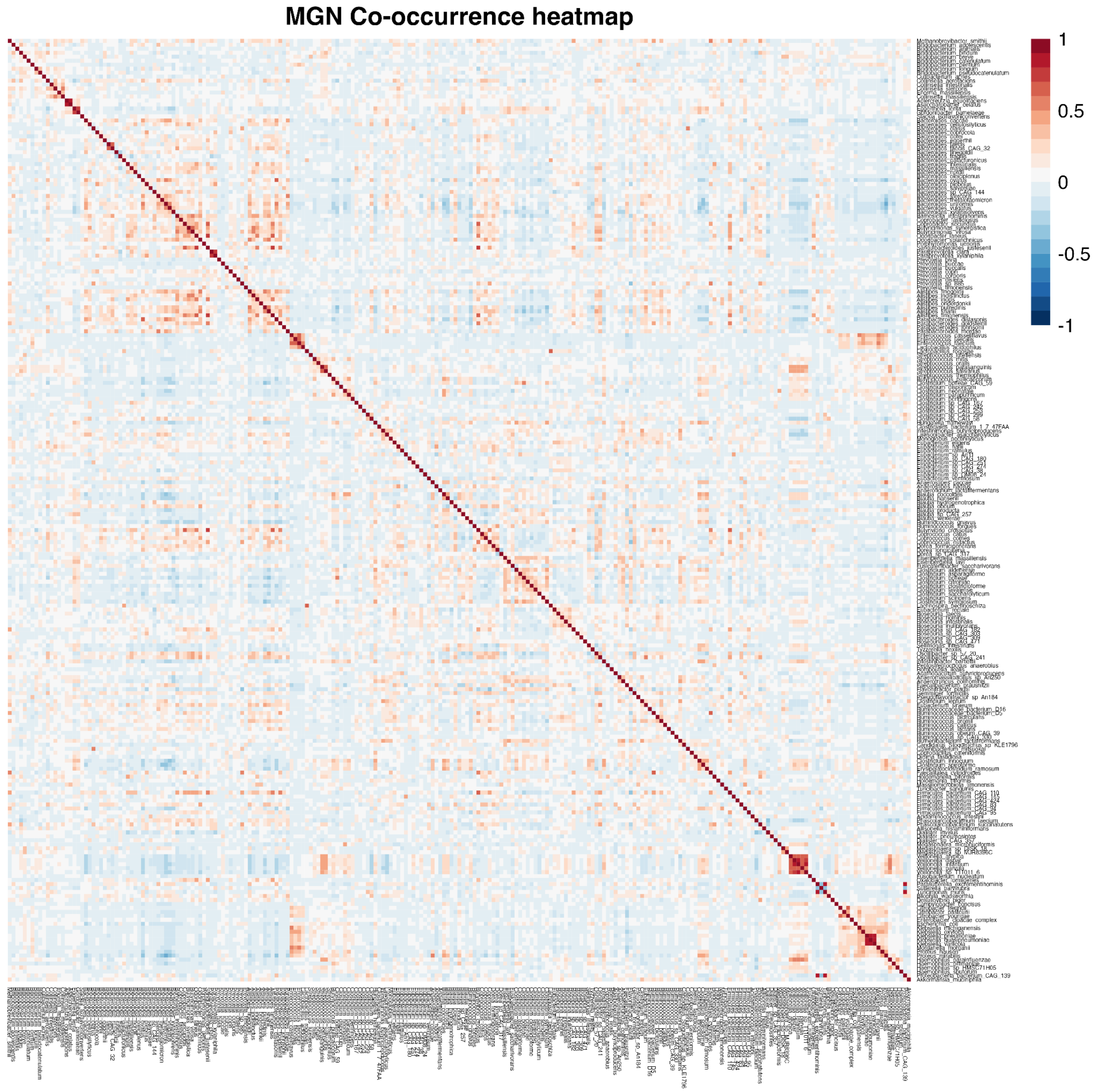


**Supplementary Figure 21:** Co-occurrence heatmap of the 237 bacteria species in the MGN dataset illustrating the Pearson correlation coefficients (r, ranging from −1 in blue to 1 in red). The co-occurrence was assessed using a Pearson correlation index matrix for the CLR-transformed compositional abundances for all 130 individuals. A complex relationship among bacteria is apparent, with 73 pairs having an $r > 0.5$, and 3 pairs with $r > 0.85$ (*Klebsiella pneumoniae - Klebsiella quasipneumoniae, Klebsiella quasipneumoniae - Turicimonas muris and Turicimonas muris - Klebsiella pneumoniae)*. Abbreviations: MGN, metagenomics; CLR, center log-ratio.

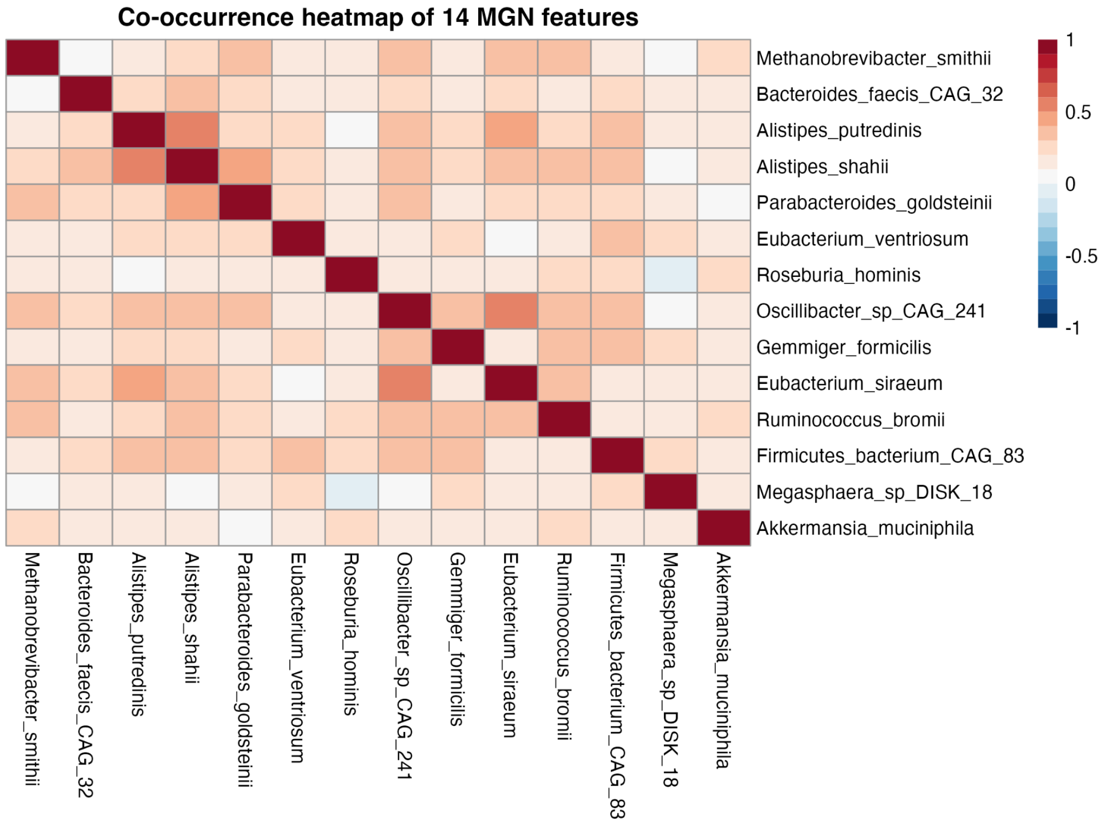


**Supplementary Figure 22:** Co-occurrence heatmap of the 14 LASSO-selected bacteria species illustrating the Pearson correlation coefficients (r, ranging from −1 in blue to 1 in red). The co-occurrence was assessed using a Pearson correlation index matrix for the CLR-transformed compositional abundances for all 130 individuals. Among the 14 features, the pairs *Eubacterium siraeum* and *Oscillibacter sp. CAG:241*, as well as *Alistipes putredinis* and *Alistipes shahii*, had the highest Pearson index of 0.5. *Roserburia hominis* and *Megasphaera sp.* DISK_18 had the lowest Pearson index of -0.03. The absence of large negative values across the heatmap suggests that these 14 features do not have mutually exclusive relationships. Abbreviations: MGN, metagenomics; LASSO, least absolute shrinkage and selection operator; CLR, center log-ratio.

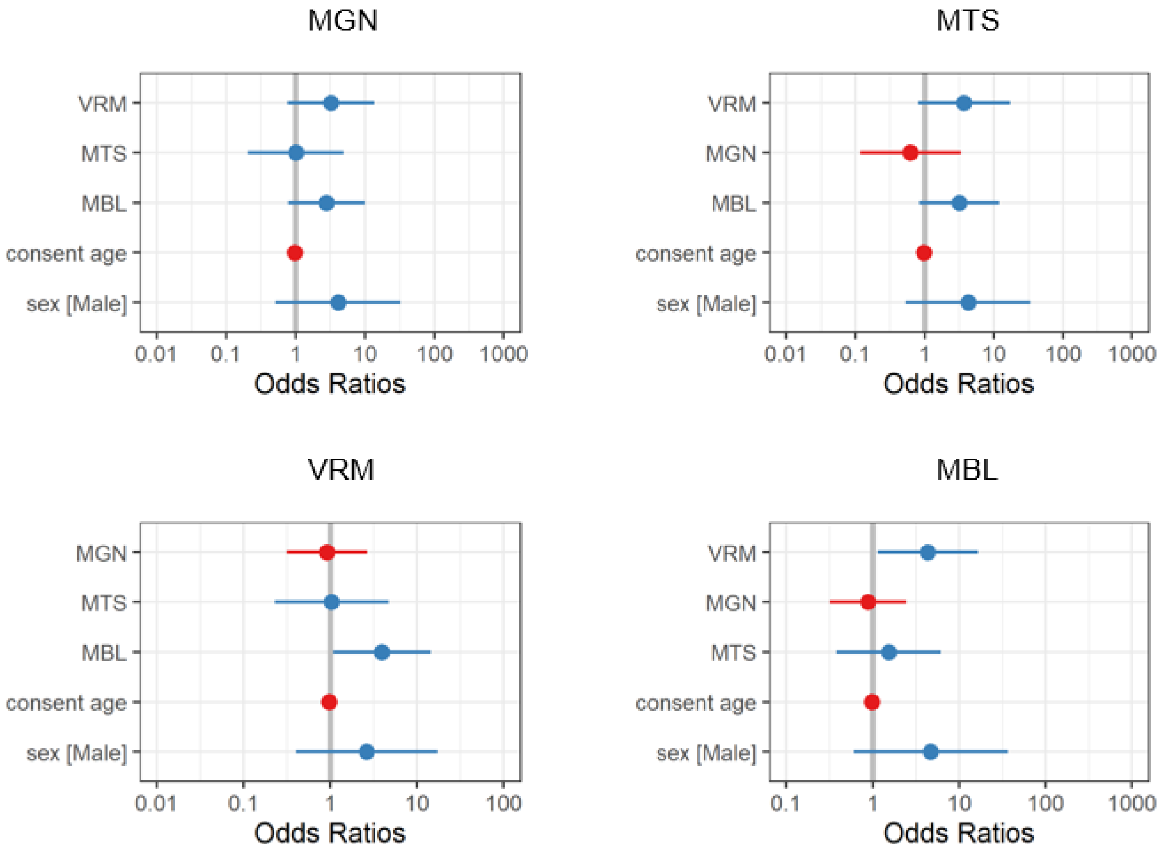


**Supplementary Figure 23**. **Leave-one-out odds ratios and 95% CIs for the combined model depict**
**ed in Figure 3.** Points represent the odds ratio for each -omic's predicted scores in a multi-omic regression where the panel title corresponds to the -omic that was removed. Lines represent 95% confidence intervals of the odds ratios. Abbreviations: MGN, metagenomics; MTS, metatranscriptomics; VRM, viromics; MBL.
